# Testis- and ovary-expressed *polo* transcripts and gene duplications affect male fertility when expressed in the germline

**DOI:** 10.1101/2024.04.05.588298

**Authors:** Paola Najera, Olivia A Dratler, Alexander B Mai, Miguel Elizarraras, Rahul Vanchinathan, Christopher A. Gonzales, Richard P. Meisel

## Abstract

Polo-like kinases (Plks) are essential for spindle attachment to the kinetochore during prophase and the subsequent dissociation after anaphase in both mitosis and meiosis. There are structural differences in the spindle apparatus between mitosis, male meiosis, and female meiosis. It is therefore possible that alleles of Plk genes could improve kinetochore attachment or dissociation in spermatogenesis or oogenesis, but not both. These opposing effects could result in sexually antagonistic selection at Plk loci. In addition, Plk genes have been independently duplicated in many different evolutionary lineages within animals. This raises the possibility that Plk gene duplication may resolve sexual conflicts over mitotic and meiotic functions. We investigated this hypothesis by comparing the evolution, gene expression, and functional effects of the single Plk gene in *Drosophila melanogaster* (*polo*) and the duplicated Plks in *Drosophila pseudoobscura* (*Dpse-polo* and *Dpse-polo-dup1*). We found that the protein-coding sequence of *Dpse-polo-dup1* is evolving significantly faster than a canonical *polo* gene across all functional domains, yet the essential structure of encoded protein appears to be retained. *Dpse-polo-dup1* is expressed primarily in testis, while other *polo* genes have broader expression profiles. Furthermore, over or ectopic expression of *polo* or *Dpse-polo* in the *D. melanogaster* male germline results in greater male infertility than ectopic expression of *Dpse-polo-dup1*. Lastly, ectopic expression of *Dpse-polo* or an ovary-derived transcript of *polo* in the male germline causes males to sire female-biased broods. However, there is no sex-bias in the progeny when *Dpse-polo-dup1* is ectopically expressed or a testis-derived transcript of *polo* is overexpressed in the *D. melanogaster* male germline. Our results therefore suggest that *Dpse-polo-dup1* may have experienced positive selection to improve its regulation of the male meiotic spindle, resolving sexual conflict over meiotic Plk functions. Alternatively, *Dpse-polo-dup1* may encode a hypomorphic Plk that has reduced deleterious effects when overexpressed in the male germline. Similarly, testis transcripts of *D. melanogaster polo* may be optimized for regulating the male meiotic spindle, and we provide evidence that the untranslated regions of the *polo* transcript may be involved in sex-specific germline functions.

## Introduction

Gametogenesis in animals is sexually dimorphic. Sex differences in gametogenesis start with the establishment of the germline, continue through meiosis, and conclude with sexually dimorphic sperm and eggs (Fuller and Spradling 2007; Whitworth *et al*. 2012; Lehtonen *et al*. 2016; Cahoon and Libuda 2019). Meiosis, a central process of gametogenesis, is highly differentiated between the sexes (Hua and Liu 2021). Male meiosis starts with a single diploid cell and produces four haploid sperm; in contrast, female meiosis produces a single haploid egg and two polar bodies from a diploid precursor (Evans and Robinson 2011; McKee *et al*. 2012). There are additional sex differences in the meiotic spindle apparatus, meiotic chromatin, chromosomal pairing, and recombination rates (Orr-Weaver 1995; McKee 1996; Sardell and Kirkpatrick 2020).

Inter-sexual differences in gametogenesis create numerous opportunities for intragenomic and intersexual conflicts (Rice 2013; Arnqvist and Rowe 2013). For example, one allele of a gene may improve some aspect of spermatogenesis, while negatively affecting oogenesis, and vice versa for the alternative allele (VanKuren and Long 2018; Hamada *et al*. 2020). This type of intralocus intersexual conflict (or sexual antagonism) may be resolved by gene duplication, followed by specialization (or subfunctionalization) of one copy for spermatogenesis or gametogenesis (Tracy *et al*. 2010; Connallon and Clark 2011; Gallach and Betrán 2011). Such germline-specific sexual subfunctionalization may be common for genes involved sex-specific or sexually dimorphic aspects of meiosis (Reis *et al*. 2011).

Intersexual conflicts likely arise because of differences between the mitotic, female meiotic, and male meiotic spindle apparatus (Orr-Weaver 1995; Savoian and Glover 2014). Despite the differences across mitotic and meiotic spindles, many genes encode proteins that are required for the mitotic, female meiotic, and male meiotic spindles. For example, the *Drosophila melanogaster* gene *mad2* encodes a protein involved in the mitotic and meiotic spindle assembly checkpoint (Li and Murray 1991; Shah and Cleveland 2000; Nicklas *et al*. 2001; Tsuchiya *et al*. 2011). In the lineage leading to *Drosophila pseudoobscura*, *mad2* was duplicated, and each copy may have evolved a specialized meiotic function in either males or females (Meisel *et al*. 2010). It is possible that sex-specific subfunctionalization of each paralog resolved an intersexual conflict that arose because of sexually dimorphic meiotic spindles.

However, there has yet to be a direct test of the hypothesis that sex differences in the meiotic spindle create sexual antagonism.

Here, we use the *Drosophila* gene *polo* as a model to explore intersexual conflicts that arise as a result of the sexually dimorphic meiotic spindle aparatus. Polo-like kinases (Plks) are essential regulators of both mitosis and meiosis across eukaryotes (Archambault and Glover 2009). Specifically, Plks are required for spindle attachment to the kinetochore during prophase and the subsequent dissociation after anaphase (Sunkel and Glover 1988; Llamazares *et al*. 1991; Donaldson *et al*. 2001). The *D. melanogaster* genome has a single Plk gene (*polo*), which is necessary for chromosome segregation during meiosis in both oogenesis and spermatogenesis (Sunkel and Glover 1988; Carmena *et al*. 1998; Herrmann *et al*. 1998; Das *et al*. 2016). Loss of function *polo* mutations affect oogenesis and early embryogenesis—from oocyte determination through meiosis and into the establishment of the embryonic sperm aster (Sunkel and Glover 1988; Tavares *et al*. 1996; Riparbelli *et al*. 2000; Mirouse *et al*. 2006). Polo is similarly required for meiotic chromosome segregation during spermatogenesis; males with *polo* mutations experience high rates of nondisjunction and produce sperm with abnormal DNA content, likely because Polo is involved in the attachment of kinetochores to the spindle apparatus (Sunkel and Glover 1988; Carmena *et al*. 1998, 2014; Herrmann *et al*. 1998). Given the differences in meiotic spindles between male and female *Drosophila* (Orr-Weaver 1995), it is possible that *polo* alleles may have sexually antagonistic effects if they improve kinetochore attachment and dissolution in spermatogenesis or oogenesis, but not both.

The location of forward (F) and reverse (R) primers used to clone cDNA from testis (poloT) and ovary (poloO) transcripts are shown. **B.** The mRNA of the alternative *polo* transcripts are diagrammed. The regions cloned from testis and ovary are shown above the polo-RB transcript.

Alternative splicing of *polo* may be one mechanism for the resolution of sexual conflict over male and female meiotic benefits. There are two polyadenylation (pA) sites within the 3’ untranslated region (UTR) of *D. melanogaster polo* (Figure 1); the two *polo* mRNA products differ in their effects on kinetochore function, pupal metamorphosis, and female fertility, possibly because of differences in translational efficiencies between transcripts with different pA sites (Llamazares *et al*. 1991; Pinto *et al*. 2011; Oliveira *et al*. 2019). Mature transcripts with the proximal pA site also have a shorter 5’-UTR (Hoskins *et al*. 2015), suggesting differences in transcription initiation sites between the mRNAs that differ in their pA site (Figure 1A). However, functional effects of the *polo* 5’-UTR have not yet been identified, and both the short and long mRNA variants encode the same protein. In addition, despite the differences between the two alternative *polo* transcripts, it is not yet known if they have sexually antagonistic effects.

**Figure 1.**
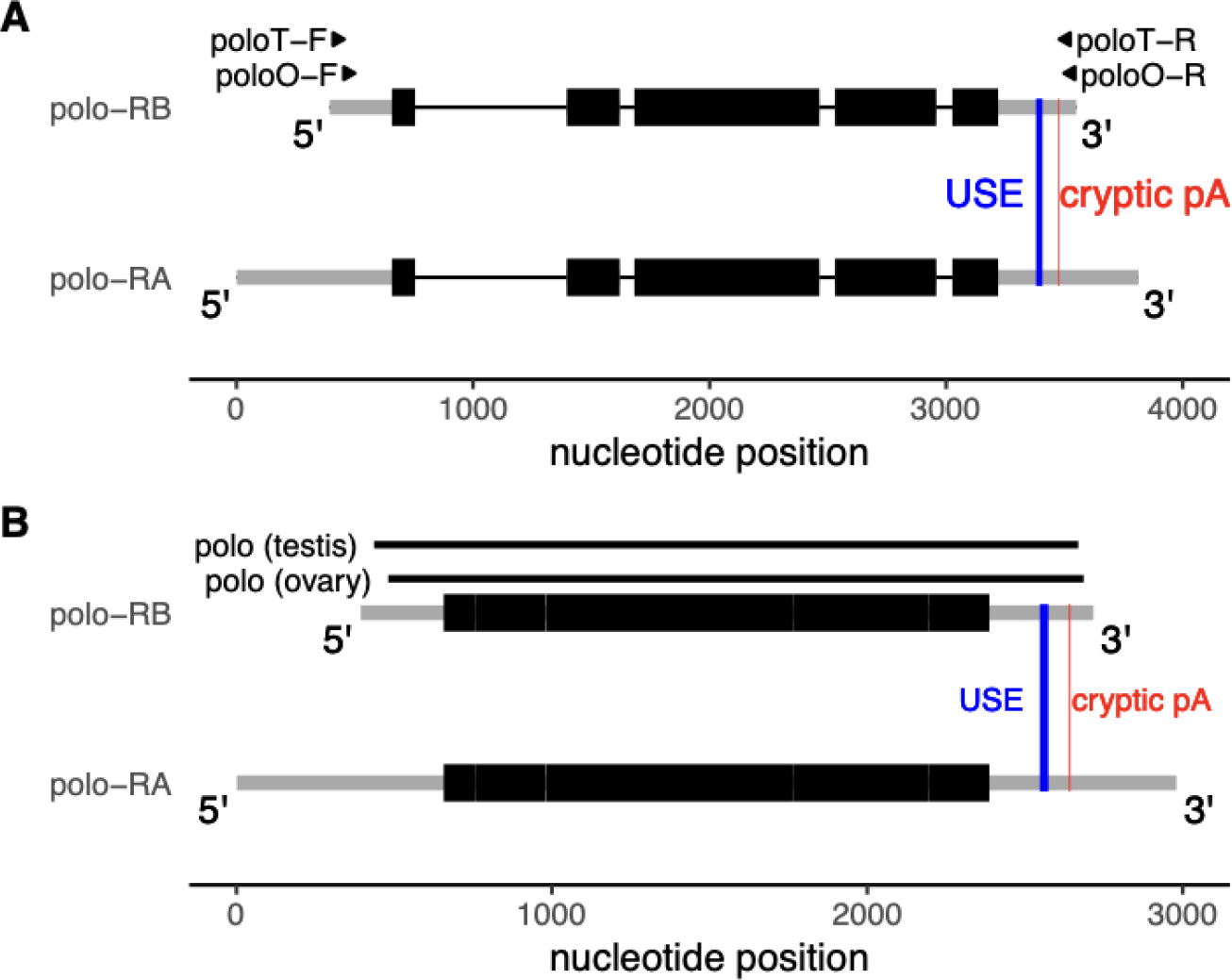
*D. melanogaster polo* is spliced into two different isoforms that differ in their untranslated regions (UTRs). The location of a conserved upstream sequence element (USE) that affects usage of the polo-RB polyadenylation (pA) site is shown by a blue line (Oliveira *et al*. 2019). A cryptic pA signal, used when the USE is mutated, is shown by a red line. **A.** The *polo* gene region is diagrammed, with two alternative transcripts of *polo* shown (polo-RA and polo-RB). Thick black boxes show protein coding exons. Medium gray bars show the 5’-UTR (left) and 3’-UTR (right). Introns are shown as thin black lines.

Gene duplication may be another way of resolving intersexual conflicts involving Plks. Plk genes have been independently duplicated multiple times during the evolution of metazoan animals, with paralogs specialized for different functions (Habedanck *et al*. 2005; Bettencourt-Dias *et al*. 2005). While *D. melanogaster* has a single Plk gene (*polo*), the *D. pseudoobscura* genome harbors two duplications (three total copies) of *polo* (Reis *et al*. 2011). In addition, *D. melanogaster polo* is autosomal (on chromosome 3L, or *Drosophila* Muller element D), but element D fused to the X chromosome in the lineage leading to *D. pseudoobscura.* Therefore, the *D. pseudoobscura* ortholog of *polo* (*Dpse-polo*) is on a neo-X chromosome. An excess of genes was duplicated from the *D. pseudoobscura* neo-X chromosome to the autosomes (Meisel *et al*. 2009), including *polo* (Reis *et al*. 2011). The same autosome independently became a neo-X chromosome in *D. willistoni*, and *mtrm* (a key interactor of *polo*) was similarly duplicated from the neo-X onto an autosome in *D. willistoni* (Xiang *et al*. 2007; Reis *et al*. 2011; Whitfield *et al*. 2013; Bonner *et al*. 2020). X-to-autosome duplications may be involved in the resolution of intersexual conflict if X-linkage is unfavorable for genes with male-specific functions, possibly because of downregulated X chromosome expression in the male germline (Betrán *et al*. 2002; Wu and Xu 2003; Emerson *et al*. 2004; Meisel *et al*. 2010). The two duplicate copies of *polo* (*polo-dup1* and *polo-dup2*) are expressed primarily in males in *Drosophila persimilis* (the sibling species of *D. pseudoobscura*), while the ancestral copy of *polo* is expressed in both sexes (Reis *et al*. 2011). The divergence in expression between *polo* paralogs is consistent with sex-specific subfunctionalization of a duplicated gene to resolve an intersexual conflict (Gallach and Betrán 2011). Notably, *polo-dup2* has a truncated protein coding sequence (it is missing the last third of the coding region), while *polo-dup1* is predicted to encode a complete Plk (Reis *et al*. 2011). This suggests *polo-dup1* may have been retained to resolve an intersexual conflict, while *polo-dup2* may be a pseudogene. Curiously, *mtrm* appears to have evolved under positive selection (Anderson *et al*. 2009), and there is evidence for divergence of Mtrm function in female meiosis across the *Drosophila* genus (Bonner and Hawley 2019). The evolutionary dynamics of *polo* and *mtrm* are therefore consistent with selection in response to intersexual conflicts, possibly as a result of sex differences in the meiotic spindle or kinetochore.

We evaluated if *polo* has sexually antagonistic effects in *Drosophila*, and we also explored if that conflict was subsequently resolved by testis-specific specialization of a *polo* gene duplication. To those ends, we examined the evolution and expression of *Dpse-polo* and a complete duplication of *polo* in the *D. pseudoobscura* genome (*Dpse-polo-dup1*). We also cloned *polo* transcripts into vectors for the GAL4>UAS binary expression system, and we determined the effect of driving their expression in the *D. melanogaster* male germline. We sampled *polo* transcripts from both the male and female *D. melanogaster* germline, in addition to *Dpse-polo* and *Dpse-polo-dup1*. We tested if expressing these different *polo* transgenes in the male germline affects male fertility and the sex ratio of progeny sired by these males.

## Materials and Methods

### Evolution of Polo protein sequences

We tested for differences in the rates of evolution of the protein sequences encoded by *Dpse-polo* and *Dpse-polo-dup1*. A previous analysis found that the nucleotide sequence of *Dpse-polo-dup1* evolves faster than *Dpse-polo* (Reis *et al*. 2011), but the rate of amino acid evolution was not directly examined. To address that shortcoming, we constructed an alignment of Dpse-Polo (XM_001353282), Dpse-Polo-dup1 (XM_002132425), and *D. melanogaster* Polo (FBtr0074839) using MUSCLE implemented in MEGA 11 for macOS with the default parameters (Edgar 2004; Stecher *et al*. 2020; Tamura *et al*. 2021). The alignment is available as Supplemental Material. We then used Tajima’s (1993) relative rate test to compare the number of amino acid substitutions in the evolutionary lineages leading to Dpse-Polo and Dpse-Polo-dup1, treating D. melanogaster Polo as the outgroup. We also compared the number of amino acid substitutions within the N-terminal serine/threonine kinase domain, the Polo box domain (PBD), the two individual Polo boxes (PB1 and PB2), and the linker between the kinase domain and PBD.

### D. pseudoobscura polo expression

We compared the expression of *Dpse-polo* and *Dpse-polo-dup1* in males and females across seven different *D. pseudoobscura* tissue samples. We first obtained normalized read counts (NRC) for all *D. pseudoobscura* genes from an RNA-seq data set in which expression was measured in four replicates from each sex for seven different tissue samples (GSE99574; Yang *et al*. 2018). We calculated the median NRC for each gene across all four replicates for each tissue-by-sex combination (NRC_TS_), and then we analyzed log_10_(NRC_TS_ + 1). We added one to each NRC_TS_ value to ensure that all values were finite (because some NRC_TS_ values were equal to zero). We compared log_10_(NRC_TS_ + 1) of *Dpse-polo* (FBgn0071596) and *Dpse-polo-dup1* (FBgn0246554) to the genome-wide distribution of log_10_(NRC_TS_ + 1) values to evaluate the relative expression of each *polo* gene in teach tissue-by-sex combination.

We used the same RNA-seq data to calculate the breadth of expression (τ) across tissues for *Dpse-polo* and *Dpse-polo-dup1* (Yanai *et al*. 2005). In our calculation of τ, we included six non-overlapping tissue samples: 1) digestive plus excretory system; 2) gonad; 3) reproductive system without gonad; 4) thorax without digestive system; 5) abdomen without digestive or reproductive system; and 6) head. We calculated τ with the following equation:

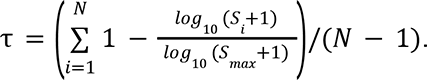

In this equation, expression of a gene in *N*=6 tissues is measured as log_10_(*S_i_* +1), where *S_i_* is the NRC_TS_ in tissue *i* for a given sex. *S_max_* is the maximum *S_i_* of the gene across all six tissue samples in a given sex. Values of τ range from 0 (equal expression in all tissues, i.e., broadly expressed) to 1 (expressed in a single tissue, i.e., narrowly expressed). We calculated τ separately for male and female tissue samples.

### Creating transgenic D. melanogaster carrying inducible polo transcripts

We cloned Plk transcripts from *D. melanogaster* testes, *D. melanogaster* ovaries, and whole *D. pseudoobscura* males. *D. melanogaster* testis and ovary tissues were dissected in Ringer’s solution from whole flies of the iso-1 strain (BDSC 2057). Ovaries and testes were dissolved overnight in TRI Reagent® on a rocker. Whole *D. pseudoobscura* males (from the MV2-25 strain) were ground in TRI Reagent® with a motorized pestle and centrifuged to remove particulates. We used the Direct-zol RNA Purification Kit (Zymo Research) to isolate RNA from each sample, following the manufacturer’s instructions.

The resultant RNA samples were used as a templates in a reverse transcription reaction (RT-PCR) with primers targeting *polo* (*D. melanogaster* testis or ovary), *Dpse-polo* (*D. pseudoobscura* males), or *Dpse-polo-dup1* (*D. pseudoobscura* males) using SuperScript™ III reverse transcriptase (Thermo Fisher Scientific). Different primer pairs were used to amplify *polo* from *D. melanogaster* ovaries (poloO) and testes (poloT) because the primers for one tissue sample would not amplify the transcript from the other tissue sample. Each of the four cDNA products was then used as a template in a PCR with the same primers and Phusion® High Fidelity DNA Polymerase (New England Biolabs). All primer pairs were located within the 5’- and 3’-UTRs of the transcripts so that they amplified the entire protein coding sequence of the respective genes (Figure 1; Supplemental Table S1). A “CACC” adapter sequence was included at the 5’ end of each forward primer to allow the PCR products to be cloned into a Gateway™ Entry vector.

We used the Gateway™ System to clone each PCR product into a vector that could be used for germline transformation of *D. melanogaster*. We first used the pENTR™/D-TOPO™ Cloning Kit to create Gateway™ Entry clones for each of the four PCR products, which we transformed into One Shot™ TOP10 Chemically Competent *Escherichia coli* cells (Thermo Fisher Scientific). We then isolated plasmids from all four cloning products with the Invitrogen™ PureLink™ Quick Plasmid Miniprep kit. We confirmed the correct insert size using PCR with the M13 primer pair. We next used the Gateway™ LR Clonase™ II Enzyme mix to recombine each of the four PCR products into the pBID-UASC-G backbone (Addgene Plasmid #35202), which contains a φC31 integrase compatible attB sequence and UAS binding sites for the GAL4 expression system (Wang *et al*. 2012). We transformed One Shot™ TOP10 Chemically Competent *E. coli* cells with each of the four recombinant plasmids. We designed primers to amplify the inserts within the pBID-UASC-G plasmid (5’-TGCCGCTGCCTTCGTTAATA-3’ and 5’-TTCCACCACTGCTCCCATTC-3’), and we confirmed that the inserts were the correct size.

We also used Sanger sequencing of the PCR products to confirm that there were no DNA sequence errors in the resulting amplifications. We finally used the Invitrogen™ PureLink™ HiPure Plasmid Filter Midiprep Kit to isolate plasmids containing each of the four PCR products.

We created transgenic *D. melanogaster* that carry one of each of the four recombinant plasmids. Each of the four plasmids was injected into *D. melanogaster* strain VK20 (BDSC 9738), which has an attP docking site at region 99F8 of chromosome 3R. All injections were performed by GenetiVision Corporation. We confirmed successful transformation via the presence of orange eyes. We balanced the third chromosome carrying each of the transgenes over a TM3,Sb chromosome. Each of these strains has the genotype UAS-poloX/TM3,Sb, where poloX refers to the specific Plk transcript (PoloO, PoloT, *Dpse-polo*, or *Dpse-polo-dup1*).

We created at least one (and no more than three) balanced strains for each transgene, with each strain originating from a different transformed founder (Supplemental Table S2).

### Assaying effects of polo transcripts on male fertility and progeny sex ratios

We tested if male germline expression of each of the four *polo* transcripts affects male fertility and sex chromosome transmission. Males with the UAS-poloX/TM3,Sb genotype were mated to females carrying a Gal4 driver construct that is expressed under the *bag of marbles* (*bam*) promoter (*P*{*bam-Gal4-VP16*}), which drives expression in the male germline (Chen and McKearin 2003; Sartain *et al*. 2011; Hart *et al*. 2018). After mating, all flies, eggs, and larvae were kept in 25x95mm vials containing cornmeal media in 25°C incubators with 12:12 light:dark cycles. Male progeny with the *P*{*bam-Gal4-VP16*}>UAS-poloX genotype were identified by wild-type bristles.

We assayed male fertility by allowing *P*{*bam-Gal4-VP16*}>UAS-poloX males to mate with wild type females from the Canton S (CanS) and Oregon R (OreR) strains. A single male and single female were combined in a 25x95mm vial with cornmeal media at 25°C, and they were observed to confirm successful copulation, as we have done previously with crosses using the same *P*{*bam-Gal4-VP16*} strain (Hart *et al*. 2018). After mating, the male was removed from the vial, and the female was allowed to lay eggs for 3–5 days at 25°C. The vials were stored at 25°C, and we counted the number of male and female progeny that emerged in each vial for 21 days after mating.

We tested for an effect of germline expression of each transgene on the number of progeny using mixed effect linear models. Our analysis compared the effects of UAS-PoloO, UAS-PoloT, UAS-*Dpse-polo*, and UAS-*Dpse-polo-dup1*. We analyzed all strains with the same transgene within a single model, treating strain as a random effect. For each comparison, we used the lme() function within the nlme package in R (Pinheiro and Bates 2000; Pinheiro *et al*. 2023) to construct a linear model with the number of progeny in a vial as a response variable, transgene as a fixed effect, and batch and strain as random effects (see Supplemental Material for R code). We tested for an effect of each transgene by separately analyzing the total number of progeny per vial, the number of male progeny, or the number of female progeny.

We also used a mixed effect logistic regression to test if the transgenes affected whether a male sires any offspring. As above, we compared the effects of UAS-PoloO, UAS-PoloT, UAS-*Dpse-polo*, and UAS-*Dpse-polo-dup1*, including all strains with the same transgene in a single model. For each comparison, we performed a logistic regression using the glmer() function in the lme4 package (Bates *et al*. 2015) to construct a model with whether a male sired progeny as a response variable (0=no, 1=yes), transgene as a fixed effect, and batch and strain as random effects (see Supplemental Material for R code). We tested for an effect of each transgene by separately analyzing if any progeny were sired, if male progeny were sired, or if female progeny were sired.

We additionally tested for differences in the sex ratio (relative numbers of male and female progeny) using mixed effect linear models. As above, we analyzed all strains with the same transgene within a single model. For each transgene, we used the lme() function in the nlme package (Pinheiro and Bates 2000; Pinheiro *et al*. 2023) to construct a linear model with the number of progeny as response variable, progeny sex (male or female) and vial as fixed effects, and batch and strain as random effects (see Supplemental Material for R code). We conclude that a transgene affects the sex ratio when progeny sex has a significant effect on the number of progeny.

## Results

### Accelerated evolution of Dpse-polo-dup1

We compared the number of amino acid substitutions in *Dpse-polo* and *Dpse-polo-dup1* to test for accelerated evolution along the lineage leading to *Dpse-polo-dup1* (Supplemental Table S3). There were significantly more amino acid substitutions in the lineage leading to *Dpse-polo-dup1* than *Dpse-polo* (χ_1_^2^=71.43, *p*<0.00001), consistent with the previously described faster evolution in the nucleotide sequence of *Dpse-polo-dup1* (Reis *et al*. 2011). Of the 567 alignable amino acid positions, 80 residues (14%) were estimated to be divergent along the lineage leading to *Dpse-polo-dup1*. In contrast, only three amino acid substitutions were identified along the lineage leading to *Dpse-polo*.

We next explored amino acid divergence along the lineage leading to *Dpse-polo* and *Dpse-polo-dup1* across the different domains of the Polo protein. Plks consist of an N-terminal serine/threonine kinase domain and a C-terminal Polo-box domain (PBD), separated by a linker. Both the kinase domain and PBD are present without any insertions or deletions in both *Dpse-polo* and *Dpse-polo-dup1*. The PBD can be further divided into Polo-box 1 (PB1) and Polo-box 2 (PB2), and there are two amino acids (histidine at position 518, and lysine at position 520) that are required to bind Polo targets (Elia *et al*. 2003a; b). Both residues are conserved in *Dpse-polo* and *Dpse-polo-dup1*. There were nine amino acids deleted in *Dpse-polo-dup1* (out of a total of 576 codons in *D. melanogaster polo*), and all nine are located in the linker (Supplemental Material). One of those amino acids was also deleted in *Dpse-polo*. Despite the structural conservation of *Dpse-polo-dup1*, there were significantly more amino acid substitutions in the kinase domain, PBD, and linker of *Dpse-polo-dup1*, relative to *Dpse-polo* (Figure 2; Supplemental Table S3). Therefore, there is a consistent signal of faster amino acid evolution in *Dpse-polo-dup1*, yet the overall structure of Polo is conserved in both *Dpse-polo* and *Dpse-polo-dup1*.

**Figure 2.**
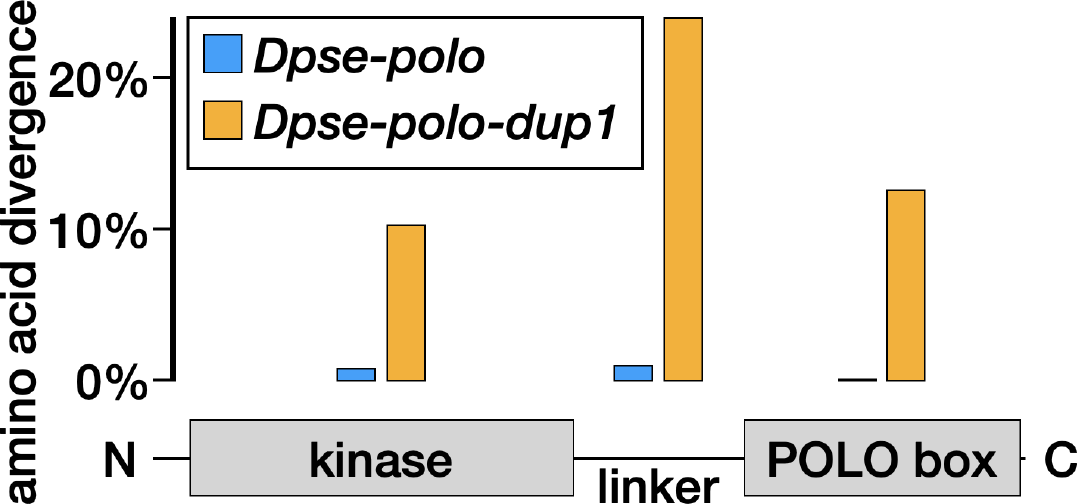
Accelerated protein coding divergence in *Dpse-polo-dup1* relative to *Dpse-polo*. Each bar shows the percent of amino acids with a substitution along the lineage leading to either *Dpse-polo* (blue) or *Dpse-polo-dup1* (orange), out of all alignable sites. Divergence was calculated within the N-terminal serine/threonine kinase domain (kinase), the C-terminal Polo-box domain (PBD), and the linker. Counts of amino acid substitutions are provided in Supplemental Table S3.

### Dpse-polo-dup1 is highly expressed in male reproductive tissues

We tested if *Dpse-polo-dup1* has male-biased expression, which would be consistent with the expression of *polo-dup1* in *D. persimilis* (Reis *et al*. 2011). To those ends, we used available RNA-seq data to compare the expression of *Dpse-polo* and *Dpse-polo-dup1* across seven different tissue samples in both males and females (Figure 3A). In each tissue sample, we observed a bimodal distribution of genome-wide expression levels, with one distribution centered close to zero (low expressed genes), and another distribution centered ∼2 orders of magnitude higher (highly expressed genes). In all sex-by-tissue combinations, *Dpse-polo* was expressed at a level within the distribution of highly expressed genes. In contrast, *Dpse-polo-dup1* was not expressed or expressed at a low level across all female tissue samples and most male samples. The notable exceptions were male samples that included reproductive tissues (whole body, reproductive system, and gonad), in which *Dpse-polo-dup1* was highly expressed, similar to *Dpse-polo*. The highest expression of *Dpse-polo-dup1* was in testis.

**Figure 3.**
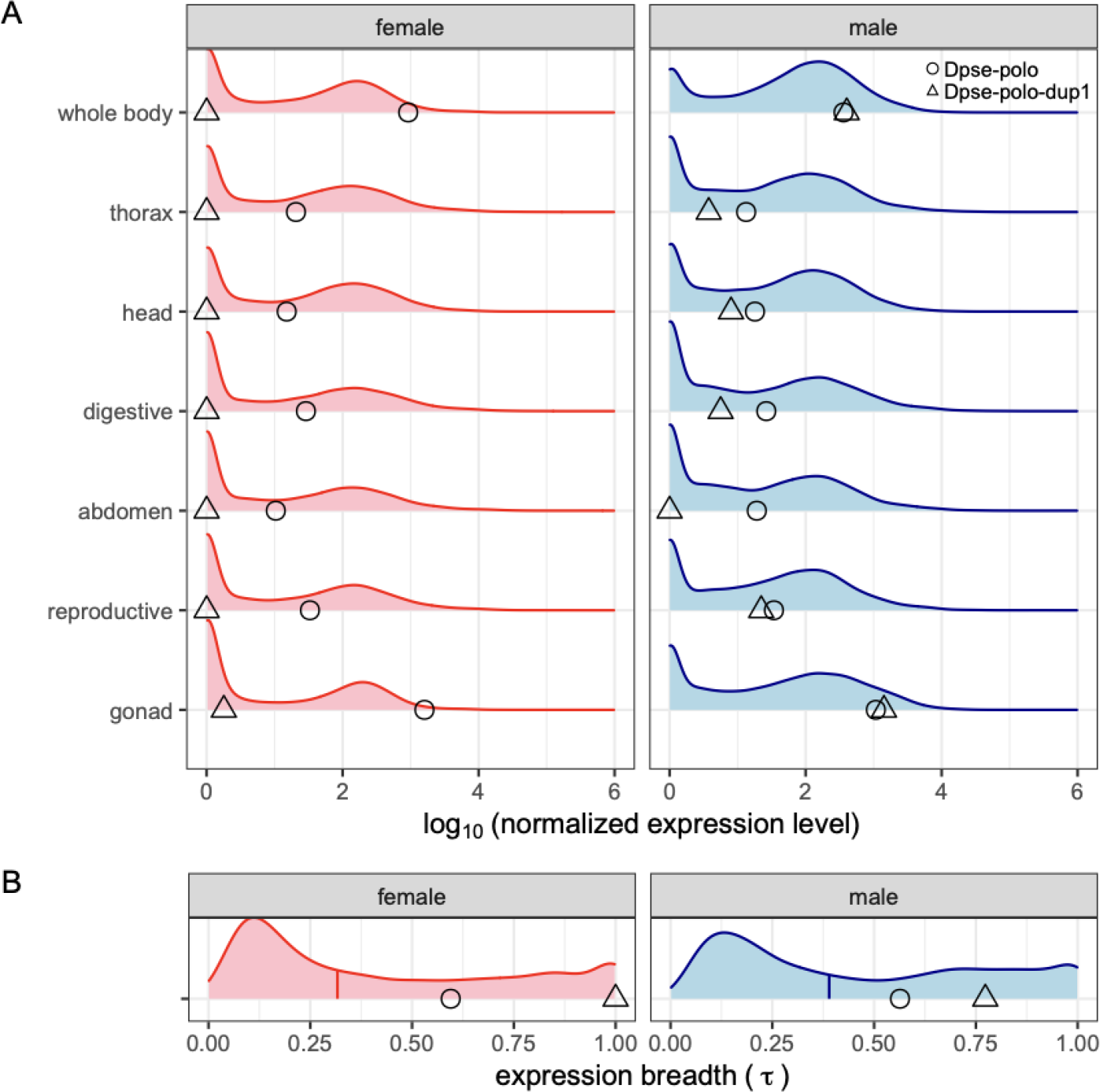
(A) Expression of *Dpse-polo* and *Dpse-polo-dup1* across seven different tissue samples in males and females. The X-axis shows the log_10_ of the median normalized expression (GSE99574; Yang *et al*. 2018). Each distribution shows the expression level of all genes in a given sex-by-tissue combination. The circles show the expression of *Dpse-polo* and the triangles show the expression level of *Dpse-polo-dup1* in each sample type. Tissue samples are whole body, thorax (with digestive system removed), head, digestive system (including excretory system), abdomen (with digestive and reproductive system removed), reproductive system (without gonad), and gonad (ovary or testis). (B) The distribution of expression breadth (τ) of all genes across six unique tissue samples (excluding whole body) in females or males is plotted. The vertical line segments within each distribution show the median value. The circles show the expression breadth of *Dpse-polo* and the triangles show the expression breadth of *Dpse-polo-dup1*.

We quantified the expression breadth of *Dpse-polo* and *Dpse-polo-dup1* using τ, which ranges from 0 (equally expressed in all tissues) to 1 (only expressed in a single tissue).

*Dpse-polo* had a similar expression breadth in both females (τ = 0.59) and males (τ = 0.56), which was larger than the median τ across the genome (Figure 3B). The high τ of *Dpse-polo* could be attributed to elevated expression in gonads relative to other tissue samples, but *Dpse-polo* was highly expressed across all tissues (Figure 3A). Surprisingly, *Dpse-polo-dup1* had the maximal τ value of 1 when expression was measured in females (Figure 3B). This is because expression was only detected in the ovary, yet *Dpse-polo-dup1* is expressed at a very low level in ovary (Figure 3A). In males, *Dpse-polo-dup1* had substantially more tissue-specific expression (τ = 0.77) than *Dpse-polo*, and this was caused by extremely high expression of *Dpse-polo-dup1* in testis (Figure 3). We therefore conclude that *Dpse-polo-dup1* has almost completely male-limited expression and strong testis-biased expression.

### Male germline expression of Dpse-polo-dup1 increases fertility

We used a GAL4>UAS system to express *Dpse-polo* and *Dpse-polo-dup1* in the *D. melanogaster* male germline. We also expressed an ovary-derived *polo* transcript (PoloO) and a testis-derived *polo* transcript (PoloT) from *D. melanogaster* (Figure 1) in the *D. melanogaster* male germline. Each of the four transcripts were cloned into a UAS expression vector to create four separate transgenes, and we refer to each as a “UAS-PoloX” transgene. We generated 1–3 transgenic strains for each of the four UAS-PoloX transgenes. Male germline expression of each transgene was under the control of a *bam*-*Gal4* driver (Chen and McKearin 2003). We mated individual bam-Gal4>UAS-PoloX males with single females, and we counted the number of male progeny and female progeny sired by each male (Supplemental Table S4).

We tested if expression of each Plk transgene in the *D. melanogaster* male germline affects the number of progeny sired. There was not a significant difference between the PoloO and PoloT transgenes on the total number of progeny, number of female progeny, or number of male progeny (all *p*>0.39; Figure 4). Similarly, there was not a significant difference between the *Dpse-polo* and *Dpse-polo-dup1* transgenes on the number of total progeny, female progeny, or male progeny (all *p*>0.24; Figure 4).

**Figure 4.**
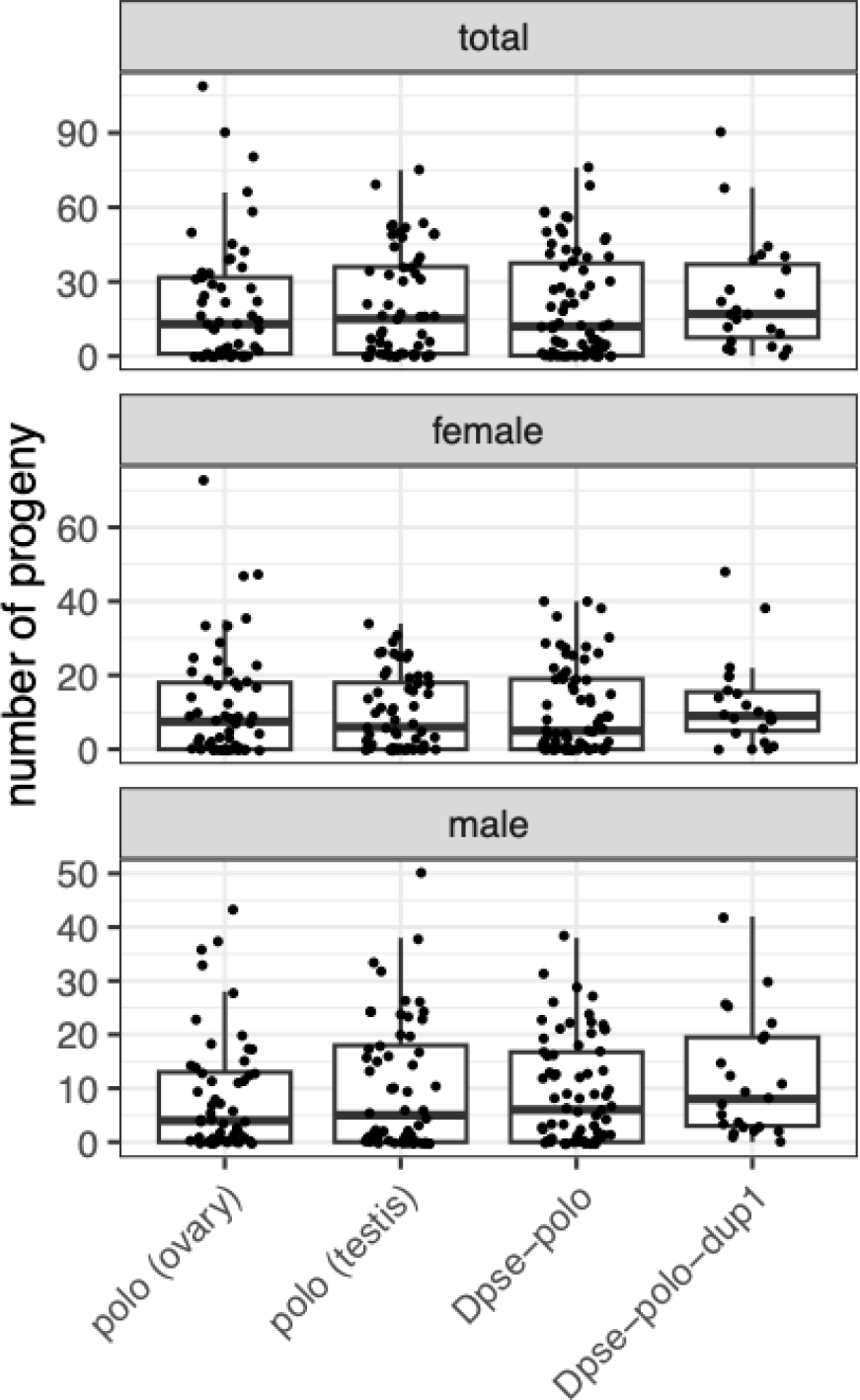
Number of progeny, number of female progeny, and number of male progeny sired by males with germline expression of different *polo* transcripts. Males carried a transgene with a *polo* transcript derived from *D. melanogaster* ovary mRNA [polo (ovary), i.e., PoloO], a *D. melanogaster* testis mRNA [polo (testis), i.e., PoloT], *Dpse-polo,* or *Dpse-polo-dup1*. Each dot shows the number of progeny sired by an individual male, and the box plots show the median and quartiles of the distribution for a given transgene.

We next tested if expressing the Plk transcripts in the *D. melanogaster* male germline affects if a male sires any progeny (i.e., whether a male sires 0 progeny or >0 progeny). Males that expressed *Dpse-polo-dup1* in their germline sired >0 progeny more frequently than males that expressed *Dpse-polo* (*z*=1.818; *p*=0.0691), PoloO (*z*=-2.108; *p=*0.0350), or PoloT (*z*=-1.673; *p*=0.0943). Approximately 20–25% of males that expressed *Dpse-polo*, PoloT, or PoloO sired 0 progeny (Supplemental Material). In contrast, only one male (out of 22, or 4.3%) who expressed *Dpse-polo-dup1* in their germline sired 0 progeny. There was not a significant difference in the number of males that sired zero progeny between those expressing *D. melanogaster* PoloO and PoloT in their germline (*z*=-1.037; *p*=0.300). We observed similar effects when we only counted male or female progeny (Supplemental Material).

### Male germline expression of ovary derived polo transcripts causes female-biased broods

We also tested if expressing different *polo* transcripts in the *D. melanogaster* male germline affects the ratio of female:male progeny sired. More female than male progeny were sired when we expressed PoloO (*F*_1,46_=9.35, *p*=0.0037) or *Dpse-polo* (*F*_1,61_=3.50, *p*=0.066) in the male germline (Figure 5). In contrast, there was not a significant difference in female and male progeny when we expressed PoloT (*F*_1,49_=0.175, *p*=0.68) or *Dpse-polo-dup1* (*F*_1,20_=0.0675, *p*=0.80) in the male germline.

**Figure 5.**
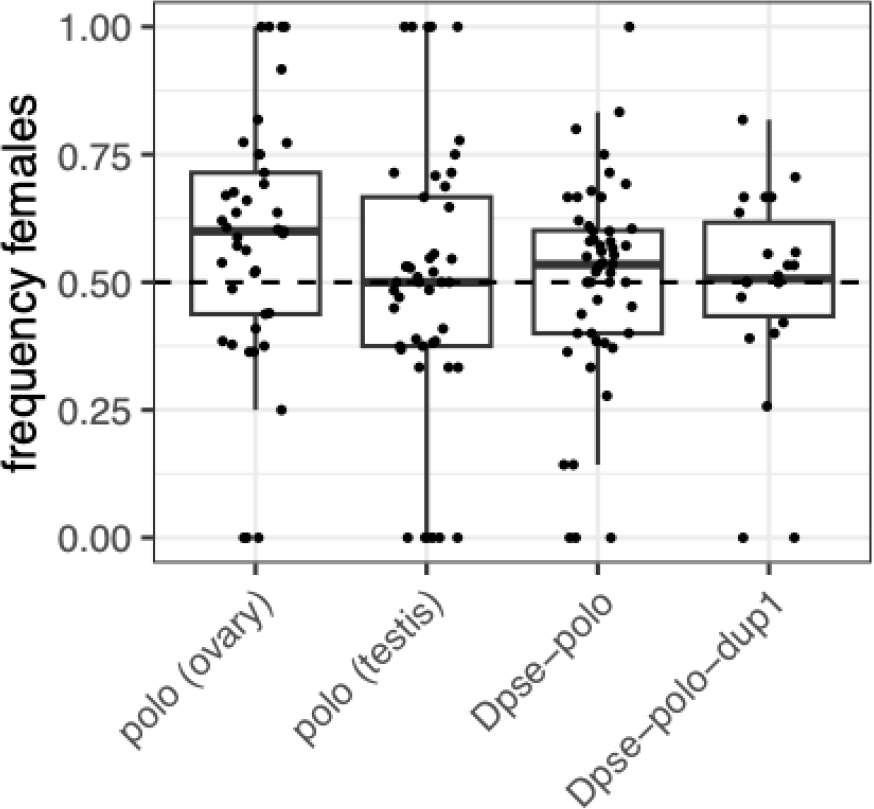
Frequency of female progeny sired by males with germline expression of different *polo* transcripts. Males carried a transgene with a *polo* transcript derived from *D. melanogaster* ovary mRNA [polo (ovary), i.e., PoloO], a *D. melanogaster* testis mRNA [polo (testis), i.e., PoloT], *Dpse-polo,* or *Dpse-polo-dup1*. Each dot shows the frequency of female progeny of progeny sired by an individual male (number of female progeny / [male + female progeny]), and the box plots show the median and quartiles of the distribution for a given transgene.

## Discussion

We showed that a duplication of *polo* in the *D. pseudoobscura* genome (*Dpse-polo-dup1*) has the conserved structure of a canonical Plk, but its amino acid sequence is fast evolving (Figure 2). We also demonstrated that *Dpse-polo-dup1* has testis-biased expression (Figure 3), suggesting specialization for male germline function. Ectopic expression of *Dpse-polo-dup1* in the *D. melanogaster* male germline increased the probability of siring progeny relative to ectopic expression of *D. melanogaster polo* or *Dpse-polo* (Figure 4), providing additional evidence that *Dpse-polo-dup1* is specialized for male germline function.

Further consistent with male germline specialization, expression of *Dpse-polo-dup1* in the *D. melanogaster* germline caused males to sire equal numbers of females and males, but male germline expression of *Dpse-polo* or a *D. melanogaster polo* transcript derived from ovary (PoloO) resulted in an excess of female progeny (Figure 5).

### Gene duplication and testis specialization of meiotic genes

Our results suggest that the rapid evolution of *Dpse-polo-dup1* may be the result of adaptive fixations of amino acid substitutions that contribute to male germline specialization. Reis et al. (2011) hypothesized that *polo* duplications with male-limited expression may accelerate male meiosis, which could provide a mechanism by which ectopic germline expression of *Dpse-polo-dup1* increases the fertility of *D. melanogaster* males. It is possible that the single copy *D. melanogaster polo* gene is constrained from germline-specific adaptation because of the diverse functions that Polo is required to perform. Plks are required for spindle attachment to and dissociation from the kinetochore in mitosis, female meiosis, and male meiosis (Sunkel and Glover 1988; Carmena *et al*. 1998; Herrmann *et al*. 1998; Archambault and Glover 2009; Das *et al*. 2016). There are functional differences between mitotic, female meiotic, and male meiotic spindles (Orr-Weaver 1995; Savoian and Glover 2014), which could create pleiotropic constraints opposing the specialization of *polo* function across mitotic and meiotic contexts (Wagner and Zhang 2011). In other words, improvements to Plk function in male meiosis could come at a cost to mitosis or female meiosis. Similar inter-sexual fitness tradeoffs have been documented in meiotic drive systems (Fishman and Saunders 2008). Duplication of *polo* in *D. pseudoobscura* may have allowed for the resolution of those pleiotropic conflicts via male meiotic specialization of *Dpse-polo-dup1* (Connallon and Clark 2011; Gallach and Betrán 2011; VanKuren and Long 2018; Hamada *et al*. 2020). Plk genes have been duplicated and subfunctionalized in other taxa (Habedanck *et al*. 2005; Bettencourt-Dias *et al*. 2005), suggesting that this may be a common mechanism to resolve pleiotropic constraints imposed by differences in the spindle apparatus across mitosis and meiosis.

If male-specific subfunctionalization of a *polo* duplication is advantageous, why does *D. melanogaster* not have subfunctionalized *polo* gene duplicates? The single *D. melanogaster polo* gene is autosomal (chromosome 3L, or Muller element D), but *Dpse-polo* became X-linked when element D fused to the X chromosome, creating a neo-X chromosome. We hypothesize that the initial retention of *Dpse-polo-dup1* was favored after *Dpse-polo* became X-linked because X chromosome expression is reduced in the male germline (Vibranovski *et al*. 2009; Meiklejohn *et al*. 2011; Wei *et al*. 2022). Reduced X expression is thought to favor the retention of autosomal duplicates of X-linked genes when those genes are required for male meiosis or spermatogenesis (Betrán *et al*. 2002; Emerson *et al*. 2004; Marques *et al*. 2005; Potrzebowski *et al*. 2008; Meisel *et al*. 2009). Therefore, X-linkage of *Dpse-polo* may have favored the initial retention of *Dpse-polo-dup1* in order to compensate for reduced expression in the male germline. This initial retention of *Dpse-polo-dup1* may have allowed for subsequent selection for male germline specialization, which resolved the antagonistic pleiotropy over meiotic and mitotic functions. This two-step process of selective retention of male germline specific paralogs could explain why *polo* duplications are not observed in other *Drosophila* species (Reis *et al*. 2011). More generally, this two-step process could explain the excess gene duplication from *Drosophila* neo-X chromosomes and rapid (possibly adaptive) evolution of testis expressed autosomal paralogs (Meisel *et al*. 2009, 2010).

### Sex ratio distortion and sexual conflict

We observed female-biased broods when we ectopically expressed either a *D. melanogaster* ovary-derived *polo* transcript (PoloO) or *Dpse-polo* in the *D. melanogaster* male germline (Figure 5). Female- or male-biased sex ratios can arise via meiotic drive or segregation distortion, and the mechanisms by which this occurs differ between oogenesis and spermatogenesis (Lindholm *et al*. 2016). In each female meiosis, only one homolog of each pair of chromosomes can segregate to the egg pole, while the remaining homolog and the sister chromatids go to polar bodies (Evans and Robinson 2011). This asymmetry in female meiosis creates an opportunity for one chromosome to outcompete its homolog to the egg pole during meiosis I, possibly because it has a “stronger” centromere (Sandler and Novitski 1957; Henikoff *et al*. 2001; Clark and Akera 2021). In contrast to oogenesis, segregation distortion in spermatogenesis typically arises when a chromosome carrying a “drive” allele is able to destroy sperm carrying an alternate allele (Larracuente and Presgraves 2012). When the drive locus is on the X chromosome (i.e., targeting Y-bearing sperm for destruction), this creates sex ratio distortion because males carrying the driving X sire an excess of daughters (e.g., Sturtevant and Dobzhansky 1936; Presgraves *et al*. 1997; Unckless *et al*. 2015).

Plks are essential for chromosome segregation during meiosis because they are required for kinetochore formation and dissociation. This role in spindle attachment to the centromere opens up the possibility for Plks to be involved in meiotic drive, possibly by affecting chromosome segregation. However, centromere drive is canonically associated with asymmetrical female meiosis, not symmetrical male meiosis (Lampson and Black 2017). It is therefore not clear how the kinetochore could affect sex chromosome transmission in the male germline, where all four meiotic chromatids are packaged into spermatids. Future work could use Plks to explore the possible associations between the male meiotic kinetochore and sex ratio distortion.

There are some similarities between the effect of *polo* on sex ratios and other known segregation distortion systems that may help us understand how *polo* expression affects sex ratios. First, most genes that cause segregation distortion in *Drosophila* are recent gene duplications that acquired germline-specific expression (Merrill *et al*. 1999; Montchamp-Moreau *et al*. 2006; Tao *et al*. 2007a; b; Helleu *et al*. 2016; Lin *et al*. 2018). For example, the *D. melanogaster Segregation Distorter* (*SD*) chromosome is preferentially transmitted relative to wild-type second chromosomes in *SD/+* heterozygous males (Temin *et al*. 1991; Larracuente and Presgraves 2012). The *Sd* locus that is responsible for *SD* drive is a truncated duplication of a gene encoding the Ran GTPase-activating protein (*RanGAP*), and the *Sd* gene is sufficient to create the driving effect of the *SD* chromosome (Merrill *et al*. 1999). In addition, simply overexpressing *RanGAP* in the male germline causes segregation distortion in a way that mimics the effect of the *SD* locus (Kusano *et al*. 2001). This driving effect of overexpression is reminiscent of the sex ratio distortion we observe when ectopically expressing PoloO or *Dpse-polo* in the male germline (Figure 5).

The mechanism by which *RanGAP* and *Sd* affect segregation distortion could be informative of how male germline *polo* expression affects progeny sex ratios. RanGAP is an integral component of nucleocytoplasmic transport (Stewart 2007), but RanGTP also affects microtubule function. For example, organization of the bipolar mitotic spindle is coordinated by Ran GTPase gradients that determine microtubule nucleation and stabilization around chromosomes (Caudron *et al*. 2005). In *Saccharomyces cerevisiae*, microtubule extension from spindle poles depends on the Ran GDP/GTP exchange factor, which functions in the opposite enzymatic direction of RanGAP (Tanaka *et al*. 2005). In addition, manipulating RanGTP in vertebrate oocytes affects meiosis II spindle assembly (Dumont *et al*. 2007), demonstrating a sex-specific meiotic effect. Genes encoding nuclear transport proteins, such as *RanGAP*, are frequently duplicated, adaptively evolving, and thought to be important loci of genetic conflict (Betrán and Long 2003; Presgraves and Stephan 2007; Presgraves 2007; Tang and Presgraves 2009; Tracy *et al*. 2010; Phadnis *et al*. 2012; Mirsalehi *et al*. 2021). The mechanisms by which RanGTP and other nuclear transport molecules affect male meiosis could be similar to how male germline expression of Plks affect sex chromosome transmission.

Second, there are multiple documented sex ratio distortion systems in *D. pseudoobscura*. One of these systems is caused by a “driving” X chromosome (known as Sex Ratio, or SR) that eliminates all Y-bearing sperm (Sturtevant and Dobzhansky 1936; Policansky and Ellison 1970). Males carrying the SR X chromosome therefore produce nearly all female offspring. In addition, male hybrids between two *D. pseudoobscura* subspecies (*pseudoobscura* and *bogotana*) are weakly fertile and sire only female offspring (Orr and Irving 2005). Decreased fertility and segregation distortion of the sex chromosomes in hybrids are both caused by incompatibilities between alleles at an X-linked locus and at least seven other interacting loci throughout the genome (Phadnis and Orr 2009; Phadnis 2011). The shared genetic architecture underlying decreased fertility and female-biased sex ratios is similar to our observation that ectopic expression of *polo* transcripts in the *D. melanogaster* male germline can affect both fertility and sex ratios. Exploring links between male fertility and sex chromosome transmission could provide further insights into how male germline expression of *polo* affects sex ratios.

Sex ratio distortion and meiotic drive are often framed as intragenomic conflicts, which are often studied independently of intralocus sexual antagonism (Lindholm *et al*. 2016). Our results provide evidence that expression of *polo* variants can create intragenomic conflict, and, more specifically, sexually antagonistic effects (Rowe *et al*. 2018). We hypothesize that *polo* alleles that optimize function in mitotic or female meiotic chromosome segregation can have deleterious effects when expressed during male meiosis. We observe these effects when we express *polo* or *Dpse-polo* in the *D. melanogaster* male germline and female-biased broods are sired. In contrast, we hypothesize that selection to optimize *Dpse-polo-dup1* for male germline function ameliorates those deleterious effects. This hypothesis explains why ectopic expression of *Dpse-polo-dup1* in the *D. melanogaster* male germline increases fertility (relative to *polo* and *Dpse-polo*) and does not skew sex ratios. Our hypothesis is also consistent with a model in which gene duplication has resolved a sexual conflict (Connallon and Clark 2011; Gallach and Betrán 2011). While our results provide a link between intralocus sexual antagonism and meiotic drive, it is not clear if sexual conflicts over meiotic functions respond to or cause meiotic drive or segregation distortion.

### Mechanisms by which polo could affect male meiosis and sex chromosome transmission

Our results are suggestive of mechanisms by which Polo transcripts could affect sex chromosome transmission and male fertility. First, we observe that ectopic expression of PoloO in the *D. melanogaster* male germline causes female-biased broods, while expression of PoloT does not (Figure 5). The PoloO and PoloT transgenes in our experiments had the same protein sequence, but they differed slightly in the UTRs they contained. PoloT has a 5’-UTR that is 45 bp longer PoloO, while PoloO has a 3’-UTR that is 17 bp longer than PoloT (Figure 1; Supplemental Table S1). It is therefore possible that a region of the 5’-UTR promotes found in PoloT equal transmission of the X and Y chromosomes, or a region of the 3’-UTR found in PoloO causes preferential transmission of the X chromosome (resulting in female-biased broods).

UTRs are known to affect both mitotic and meiotic functions of Plks. For example, the two *polo* transcripts in *D. melanogaster* differ in the lengths of their 3’-UTRs (Figure 1), which affects translational efficiency and possibly kinetochore function, metamorphosis, and female fertility (Llamazares *et al*. 1991; Pinto *et al*. 2011; Oliveira *et al*. 2019). In addition, an allele in the human PLK1 3’-UTR affects mRNA secondary structure and stability (Akdeli et al. 2014), and shorter 3’-UTRs in many genes are associated with enhanced cell proliferation (Sandberg *et al*. 2008; Mayr and Bartel 2009). Some of these effects are caused by different pA sites, which should not differ between PoloO and PoloT—they share the same pA site that was engineered into their cloning backbone (Wang *et al*. 2012). However, the transcription rate of *polo* affects pA site selection, possibly via auto-regulatory feedback (Pinto *et al*. 2011). The GAL4>UAS system that we used may therefore have affected expression levels of *polo* transcripts in a way that shifted the relative usage of the pA site in the cloning backbone and a cryptic pA site in the 3’-UTR (Figure 1). It is also possible that the additional sequence in the PoloO 3’-UTR may affect the testis function of *polo* via effects on transcript stability or translational efficiency.

It is notable that a transcript that appears to affect X chromosome transmission in spermatogenesis was cloned from the ovary (PoloO), whereas a testis-derived transcript (PoloT) has no such effects (Figure 5). We cloned different transcripts from ovary and testis because the PCR primers that amplified *polo* transcripts in one tissue sample did not work in the other tissue sample. Upon examination of RNA-seq reads mapped to the *polo* locus in *D. melanogaster*, we discovered that the testis and ovary transcripts of *polo* may have atypical UTR configurations (Supplemental Figure S1). Specifically, testis transcripts appear to have the longer 5’-UTR (similar to polo-RA) and the shorter 3’-UTR (similar to polo-RB). In contrast, ovary transcripts appear to have the shorter 5’-UTR (similar to polo-RB) and the longer 3’-UTR (similar to polo-RA). These different UTR configurations may explain why we could amplify a longer 5’-UTR in PoloT and a longer 3’-UTR in PoloO (Figure 1). These differences are also consistent with our hypothesis that a sequence in the 5’-UTR of *polo* has testis-beneficial effects or a sequence in the 3’-UTR is ovary-beneficial. These testis- and/or ovary-specific effects may provide a mechanism for sexual conflict over transcript expression or splicing, possibly via transcript stability or translational efficiency.

A second important observation is that expression of *polo* or *Dpse-pol*o in the *D. melanogaster* male germline decreases male fertility relative to *Dpse-polo-dup1* (Figure 4). We hypothesized that the higher relative fertility of males expressing *Dpse-polo-dup1* is caused by amino acid substitutions that optimize the protein for testis function. An alternative hypothesis is that high testis expression of Plks in the male germline decreases fertility, and *Dpse-polo-dup1* encodes a Plk with a mild loss of function with a lower fertility cost. In this hypothesis, ectopic expression of *Dpse-polo-dup1* would be less costly than expression of the fully functional Plks encoded by *polo* and *Dpse-polo*. Negative effects of high *polo* expression have been shown in *D. melanogaster* intestinal stem cells, where constitutively active Polo suppresses intestinal stem cell proliferation, induces abnormal accumulation of β-tubulin in cells, and drives stem cell loss via apoptosis (Zhang *et al*. 2023). However, other experiments have shown Polo overexpression by 2.5-fold using GAL4>UAS does not affect its physiological function in mitosis (Martins *et al*. 2009). It therefore remains to be determined *Dpse-polo-dup1* has fewer negative effects when ectopic expressed in the male germline, or if it has beneficial effects because of selection for testis specialization.

We hypothesize that changes to *polo* transcript stability, translational efficiency, or protein coding sequence affect sex chromosome segregation in male meiosis. This hypothesis is motivated by the observation that mutations to *polo* cause high rates of nondisjunction and sperm with abnormal DNA content (Sunkel and Glover 1988; Carmena *et al*. 1998, 2014; Herrmann *et al*. 1998), and we observed that ectopic expression of transcripts in the *D. melanogaster* male germline affected the sex ratio of broods in our experiments (Figure 5). Polo may affect chromosomal transmission through its interactions with Mei-S332. Mei-S332 associates with centromeres in prometaphase of meiosis I, and phosphorylation by Polo is required for removal of Mei-S332 during segregation of sister chromatids in anaphase II (Goldstein 1981; Kerrebrock *et al*. 1992; Tang *et al*. 1998; Clarke *et al*. 2005). Mutation of *mei-S332* causes nondisjunction during meiosis II because of defective sister chromosome cohesion after metaphase I, which affects orientation going into meiosis II (Davis 1971; Goldstein 1980). Nondisjunction of autosomes could decrease fertility by increasing the frequency of autosomal aneuploids. Another outcome of elevated meiosis II nondisjunction is that *mei-S332* mutant males produce an excess of XX sperm (i.e., co-inheritance of sister chromatids), relative to XY sperm (co-inheritance of homologous chromatids), in addition to an excess of nullo-XY sperm (Kerrebrock *et al*. 1992). These two predictions could provide a functional link between sex ratio distortion and male fertility. However, while an excess of XX sperm would cause males to sire female-biased broods, an excess of nullo-XY sperm would lead to male-biased broods, effectively canceling each other out. If *polo* expression affects Mei-S332 function, it is therefore not clear how this would result in female-biased broods.

## Conclusions

We showed that a fast evolving, testis-expressed duplication of *polo* in the *D. pseudoobscura* genome (*Dpse-polo-dup1*) does not impose fertility costs nor does it skew progeny sex ratios when expressed in the *D. melanogaster* male germline. In contrast, ectopic testis expression of ovary derived *polo* genes and transcripts causes males to sire female-biased broods. These results are consistent with adaptive specialization of *Dpse-polo-dup1* for male germline-specific function, possibly related to unique requirements associated with the male meiotic spindle apparatus. Alternatively, *Dpse-polo-dup1* may be a hypomorphic Plk variant that does not have deleterious effects when over-expressed in the male germline, in contrast to other Plks. The initial duplication of *polo* may have been selectively retained because neo-X-linkage caused decreased male germline expression of the ancestral *polo* locus, favoring an autosomal paralog to compensate. This could explain why *D. pseudoobscura* has a testis-expressed paralog of *polo* but *D. melanogaster* does not. These results provide evidence for divergent selection pressures on spindle assembly genes in mitosis, female meiosis, and male meiosis. We hypothesize that these divergent selection pressures create pleiotropic conflicts or sexual antagonism, which can then be resolved by duplication and germline-specific specialization of a paralog.

## Supporting information

Code for analyzing RNA-seq data

RNA-seq data

Code for analyzing overexpression experiments

fertility data from overexpression experiments

alignment of polo protein coding sequences

## Acknowledgements

We thank Samantha Pacheco and Taylor Nunley for assistance with experiments, and other members of the Meisel lab for valuable discussions. This work was supported by startup funds from the University of Houston to RPM and a University of Houston Summer Undergraduate Research Fellowship to RV.

## Supplemental Figures

**Supplemental Figure S1.**
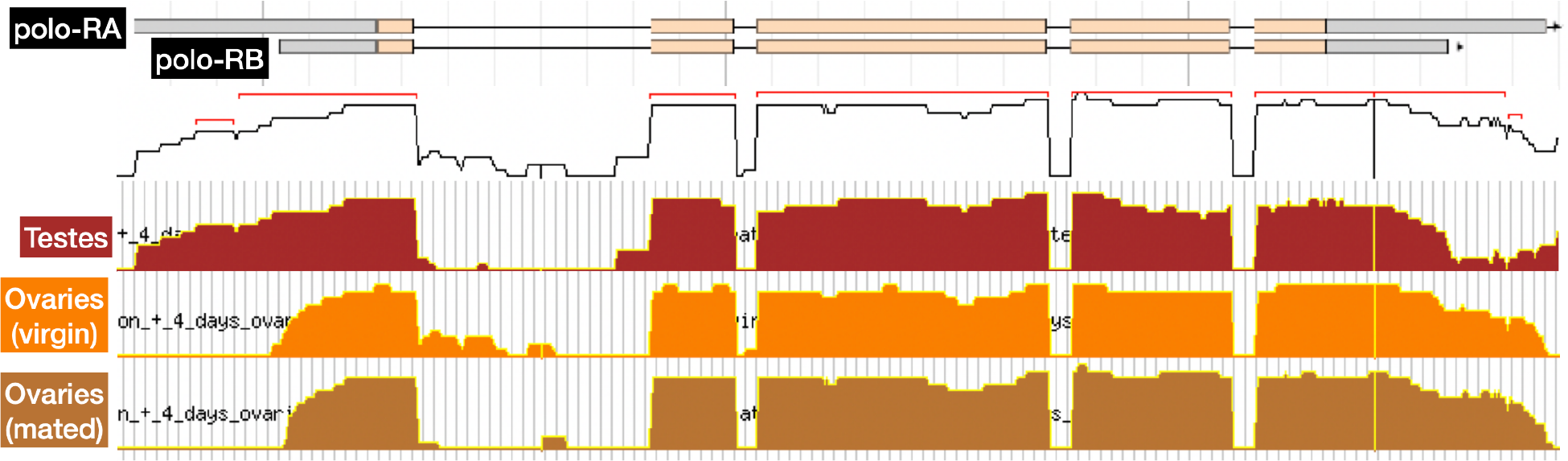
RNA-seq read mapping across the *polo* gene region. Transcript structures of polo-RA and polo-RB are shown, with UTRs in gray and protein coding sequence in tan. RNA-seq reads mapped per nucleotide position are shown for samples from testes, ovaries (virgin females), and ovaries (mated females). Average coverage across all samples is shown as a line graph, with predicted exons indicated by red brackets. Figure downloaded and modified from FlyBase JBrowse. Original RNA-seq data from modENCODE (Brown *et al*. 2014).

