## Supplementary material for "Testis- and ovary-expressed *polo* transcripts and gene duplications affect male fertility when expressed in the germline": Code for analyzing RNA-seq data

### RNA-seq Data Analysis

2024-04-05

#### Load data and setup initial objects

Load the *Drosophila pseudoobscura* RNA-seq data:

```
DpseNRCfb <- read.delim("dpse.nrc.FB.txt")
```

The following sample types are present in these data:

| sample_ID | sample_type |
|---|---|
| ac | abdomen without digestive or reproductive system |
| dg | digestive plus excretory system |
| go | gonad |
| hd | head |
| re | reproductive system without gonad |
| tx | thorax without digestive system |
| wb | whole body |

Male (`m`) and female (`f`) samples were collected for each sample type. There are four replicates per sample type. Calculate the `median` expression value across all replicates for each sex by sample type combination:

```
DpseAvg <- data.frame(
    FBgnID = DpseNRCfb$FBgnID,
    dpse_f_ac = apply(DpseNRCfb[, 2:5 ],1, median),
    dpse_f_dg = apply(DpseNRCfb[, 6:9 ],1, median),
    dpse_f_go = apply(DpseNRCfb[,10:13],1, median),
    dpse_f_hd = apply(DpseNRCfb[,14:17],1, median),
    dpse_f_re = apply(DpseNRCfb[,18:21],1, median),
    dpse_f_tx = apply(DpseNRCfb[,22:25],1, median),
    dpse_f_wb = apply(DpseNRCfb[,26:29],1, median),
    dpse_m_ac = apply(DpseNRCfb[,30:33],1, median),
    dpse_m_dg = apply(DpseNRCfb[,34:37],1, median),
    dpse_m_go = apply(DpseNRCfb[,38:41],1, median),
    dpse_m_hd = apply(DpseNRCfb[,42:45],1, median),
    dpse_m_re = apply(DpseNRCfb[,46:49],1, median),
    dpse_m_tx = apply(DpseNRCfb[,50:53],1, median),
    dpse_m_wb = apply(DpseNRCfb[,54:57],1, median)
)
```

The two *D. pseudoobscura polo* genes (*Dpse-polo* and *Dpse-polo-dup1*) have the following annotation IDs:

| | Dpse_polo | Dpse_polo_dup1 |
|---|---|---|
| ID | GA11545 | GA25172 |
| transcript | XM_001353282.5 | XM_002132425.3 |
| geneID | LOC4812761 | LOC6903594 |
| protein | XP_001353318.5 | XP_002132461.2 |
| FlyBaseID | FBgn0071596 | FBgn0246554 |

#### Compare expression of D. pseudoobscura polo genes versus genome-wide distribution

Generate a ridgeplot comparing *Dpse-polo* and *Dpse-polo-dup1* expression against the distribution of expression in each sex by sample type combination:

```
library(ggplot2)
library(ggridges)
library(cowplot)
library(grid)

data.framer <- function(FBIDs, Sex, Tissue, Expr){
    data.frame(
        FBgnID = FBIDs,
        sex = Sex,
        tissue = Tissue,
        express = Expr
    )
}
DpseAvg.df <- rbind(
    data.framer(DpseAvg$FBgnID, "female","abdomen", DpseAvg$dpse_f_ac),
    data.framer(DpseAvg$FBgnID, "female","digestive", DpseAvg$dpse_f_dg),
    data.framer(DpseAvg$FBgnID, "female"," gonad", DpseAvg$dpse_f_go),
    data.framer(DpseAvg$FBgnID, "female","head", DpseAvg$dpse_f_hd),
    data.framer(DpseAvg$FBgnID, "female"," reproductive", DpseAvg$dpse_f_re),
    data.framer(DpseAvg$FBgnID, "female","thorax", DpseAvg$dpse_f_tx),
    data.framer(DpseAvg$FBgnID, "female","whole body", DpseAvg$dpse_f_wb),
    data.framer(DpseAvg$FBgnID, "male","abdomen", DpseAvg$dpse_m_ac),
    data.framer(DpseAvg$FBgnID, "male","digestive", DpseAvg$dpse_m_dg),
    data.framer(DpseAvg$FBgnID, "male"," gonad", DpseAvg$dpse_m_go),
    data.framer(DpseAvg$FBgnID, "male","head", DpseAvg$dpse_m_hd),
    data.framer(DpseAvg$FBgnID, "male"," reproductive", DpseAvg$dpse_m_re),
    data.framer(DpseAvg$FBgnID, "male","thorax", DpseAvg$dpse_m_tx),
    data.framer(DpseAvg$FBgnID, "male","whole body", DpseAvg$dpse_m_wb)
)

DpseRNAseq <- ggplot(DpseAvg.df, aes(y=tissue, x=log10(express+1), fill=sex, color=sex)) +
    geom_density_ridges(scale=0.9) +
    scale_color_manual(values=c("female"="red", "male"="darkblue")) +
    scale_fill_manual(values=c("female"="pink", "male"="lightblue")) +
    geom_point(data=subset(DpseAvg.df, FBgnID=="FBgn0071596"),
               color="black", fill="black", shape=1, size=3) +
    geom_point(data=subset(DpseAvg.df, FBgnID=="FBgn0246554"),
               color="black", fill="black", shape=2, size=3) +
```

```
    scale_x_continuous(expression(log[10]~"(normalized"~expression~"level)"),
                       limits=c(0, 6)) +
    scale_y_discrete("") +
    facet_wrap(~sex, ncol=2) +
    theme_bw() +
    theme(legend.position = "none")

ggdraw(DpseRNAseq) +
  draw_label("\u25cb", size=11, x=0.84, y=0.9) +
  draw_label("\u25b3", size=11, x=0.84, y=0.87) +
  draw_label("Dpse-polo", size=8, x=0.85, y=0.9, hjust=0) +
  draw_label("Dpse-polo-dup1", size=8, x=0.85, y=0.87, hjust=0)
```

```
## Picking joint bandwidth of 0.145
```

```
## Picking joint bandwidth of 0.138
```

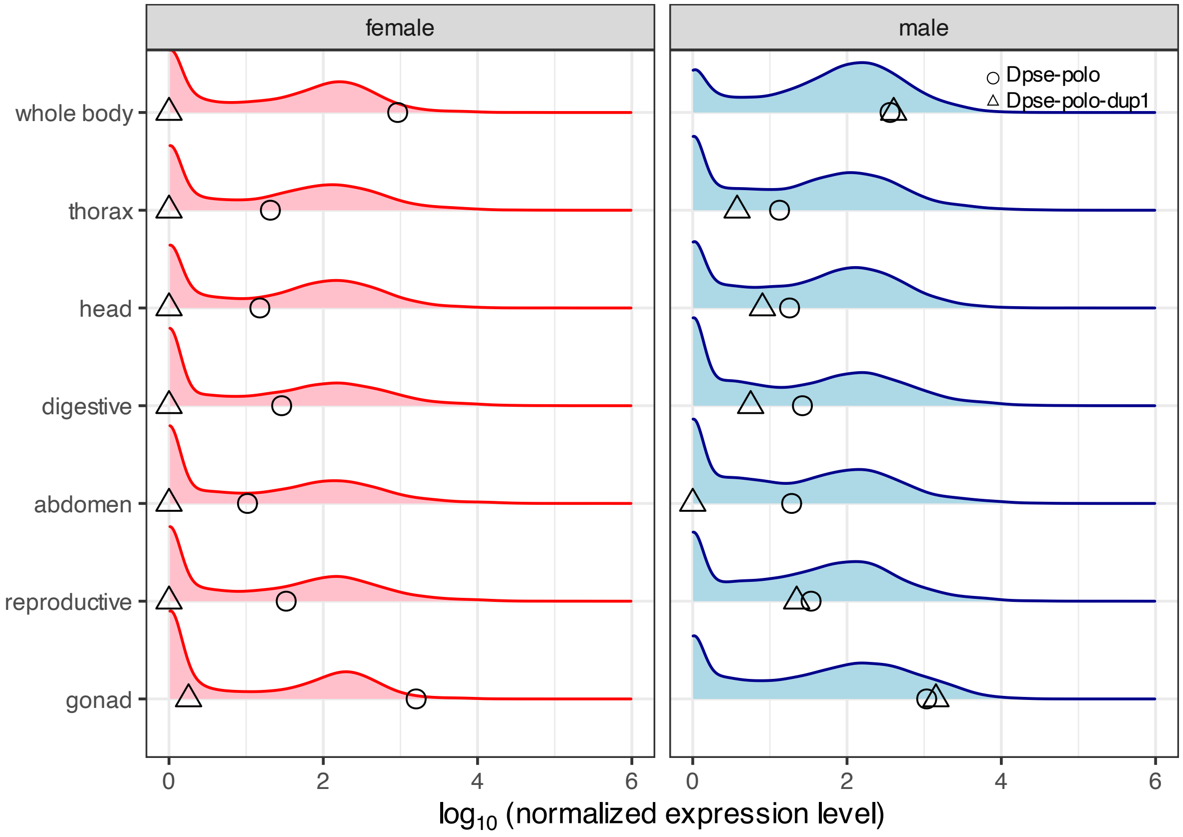

#### Compare tissue-specificity (tau) of D. pseudoobscura polo genes against genome

Calculate tau using the equation:

$$\tau = \sum_{i=1}^{N} \frac{1 - \frac{\log S_i}{\log S_{max}}}{N-1}$$

For our data, N = 6 (see tissue list above). We will define a function to calculate tau:

```
tau.calc <- function(datas){
    N <- length(datas)
    Smax <- max(datas)
    taus <- 1 - log10(datas+1)/log10(Smax+1)
    sum(taus)/(N-1)
}
DpseAvg$tau_female <- apply(
  subset(DpseAvg, select=c(dpse_f_ac, dpse_f_dg, dpse_f_go, dpse_f_hd, dpse_f_re, dpse_f_tx)),
  1, tau.calc)
DpseAvg$tau_male <- apply(
  subset(DpseAvg, select=c(dpse_m_ac, dpse_m_dg, dpse_m_go, dpse_m_hd, dpse_m_re, dpse_m_tx)),
  1, tau.calc)
```

Prepare object to be used to make graphs showing tau values and plot:

```
YaxisSpacer = "                          "
DpseAvgTau <- na.omit(rbind(
    data.frame(gene=DpseAvg$FBgnID, tau=DpseAvg$tau_female, sex="female", Yaxis=YaxisSpacer),
    data.frame(gene=DpseAvg$FBgnID, tau=DpseAvg$tau_male, sex="male", Yaxis=YaxisSpacer)))

DpseTau <- ggplot(DpseAvgTau, aes(y=Yaxis, x=tau, fill=sex, color=sex)) +
    geom_density_ridges(scale=0.9, quantile_lines = TRUE, quantiles = 2) +
    scale_color_manual(values=c("female"="red", "male"="darkblue")) +
    scale_fill_manual(values=c("female"="pink", "male"="lightblue")) +
    geom_point(data=subset(DpseAvgTau, gene=="FBgn0071596"),
               color="black", fill="black", shape=1, size=3) +
    geom_point(data=subset(DpseAvgTau, gene=="FBgn0246554"),
               color="black", fill="black", shape=2, size=3) +
    scale_x_continuous(expression("expression"~breadth~"("~tau~")"), limits=c(0,1)) +
    scale_y_discrete("") +
    facet_wrap(~sex, ncol=2) +
    theme_bw() +
    theme(legend.position = "none")

ggdraw(DpseTau) +
  draw_label("\u25cb", size=11, x=0.84, y=0.9) +
  draw_label("\u25b3", size=11, x=0.84, y=0.87) +
  draw_label("Dpse-polo", size=8, x=0.85, y=0.9, hjust=0) +
  draw_label("Dpse-polo-dup1", size=8, x=0.85, y=0.87, hjust=0)
```

```
## Picking joint bandwidth of 0.0441
```

```
## Picking joint bandwidth of 0.0417
```

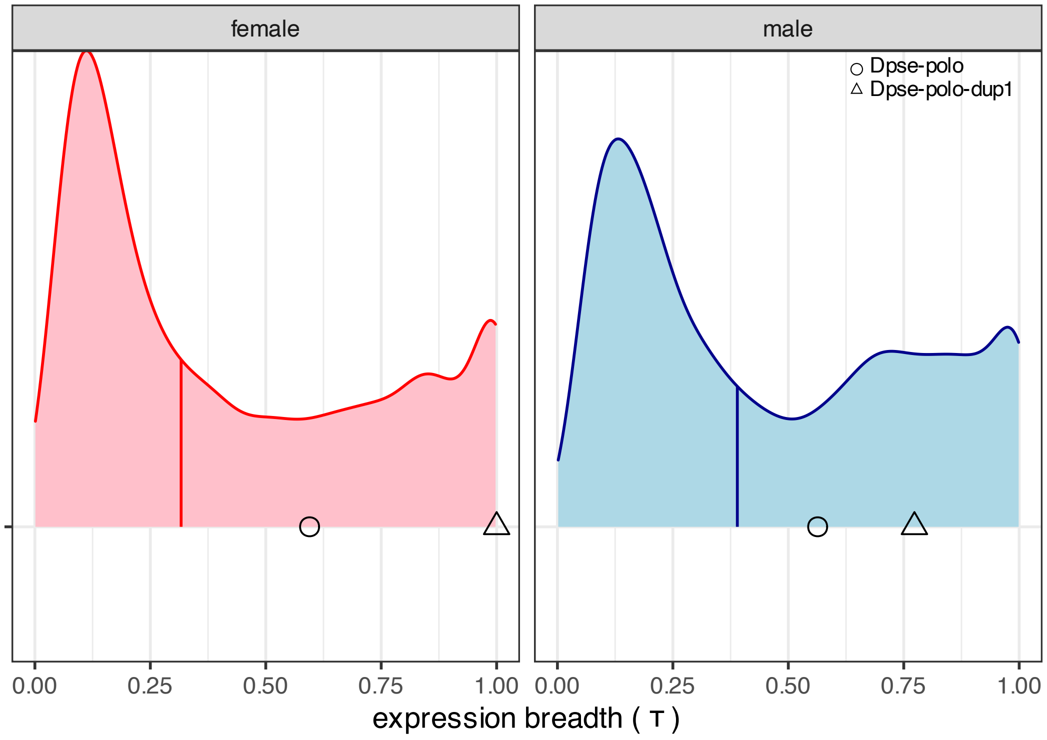
female
male
Dpse-polo
Dpse-polo-dup1
0.00
0.25
0.50
0.75
1.00
0.00
0.25
0.50
0.75
1.00
expression breadth ( τ )
