## Supplementary material for "Testis- and ovary-expressed *polo* transcripts and gene duplications affect male fertility when expressed in the germline": Code for analyzing overexpression experiments

### How do polo transgenes affect male fertility and progeny sex ratios?

2023-04-08

#### Prepare data and load packages for analysis

4-GA11545 corresponds to *Dpse-polo*.

5-GA25172 corresponds to *Dpse-polo-dup1*.

https://www.ncbi.nlm.nih.gov/pmc/articles/PMC3053365/

##### Load required packages

```
library(tidyr)
library(ggplot2)
library(nlme) # for lme to do mixed effect model
library(lme4) # for glmer to do logistic regression with random effects
```

##### Load the data

```
bams <- read.delim("bam-polo.tsv")
```

#### What is the effect of each transgene on the total number of progeny?

##### Prepare data for analysis

```
bams$progeny <- bams$males + bams$females
bamCounts <- data.frame(
    date = bams$Cross.date,
    vial = bams$vial,
    strain = bams$strain,
    counts = c(bams$males, bams$females, bams$progeny),
    sex = c(
        rep("male",length(bams$males)),
        rep("female",length(bams$females)),
        rep(" total",length(bams$progeny)))
)
bamCounts <- bamCounts %>%
  separate(strain, into=c("transgene", "strainID"), sep=" ", remove=FALSE)
```

##### Create function to performed linear mixed effects model

```
nlme_count_strain <- function(crossData){
    anova(
        lme(fixed = counts ~ transgene,
            data = crossData,
            random = list(~1|date, ~1|strainID)
        )
    )
}
```

#### Test effect on the total number of progeny

##### Do Dpse-polo and Dpse-polo-dup1 differentially affect the total number of progeny?

```
nlme_count_strain(
  subset(bamCounts, sex==" total"
        & (transgene=="4-GA11545" | transgene=="5-GA25172")))
```

```
##             numDF denDF F-value p-value
## (Intercept)     1    83 9.80123  0.0024
## transgene       1    83 0.85944  0.3566
```

##### Do Dmel PoloO and PoloT differentially affect the total number of progeny?

```
nlme_count_strain(
  subset(bamCounts, sex==" total"
        & (transgene=="6-PoloT" | transgene=="7-PoloO")))
```

```
##             numDF denDF   F-value p-value
## (Intercept)     1    95 15.880085  0.0001
## transgene       1    11  0.049549  0.8279
```

#### Test effect on the number of female progeny

##### Do Dpse-polo and Dpse-polo-dup1 differentially affect the number of female progeny?

```
nlme_count_strain(
  subset(bamCounts, sex=="female" & (transgene=="4-GA11545" | transgene=="5-GA25172")))
```

```
##             numDF denDF  F-value p-value
## (Intercept)     1    83 9.904780  0.0023
## transgene       1    83 0.445245  0.5065
```

##### Do Dmel PoloO and PoloT differentially affect the number of female progeny?

```
nlme_count_strain(
  subset(bamCounts, sex=="female"
        & (transgene=="6-PoloT" | transgene=="7-PoloO")))
```

```
##             numDF denDF   F-value p-value
## (Intercept)     1    95 14.577479  0.0002
## transgene       1    11  0.782888  0.3952
```

#### Test effect on the number of male progeny

##### Do Dpse-polo and Dpse-polo-dup1 differentially affect the number of male progeny?

```
nlme_count_strain(
  subset(bamCounts, sex=="male"
        & (transgene=="4-GA11545" | transgene=="5-GA25172")))
```

```
##              numDF denDF  F-value p-value
## (Intercept)     1    83 9.351115  0.0030
## transgene       1    83 1.389398  0.2419
```

##### Do Dmel PoloO and PoloT differentially affect the number of male progeny?

```
nlme_count_strain(
  subset(bamCounts, sex=="male"
        & (transgene=="6-PoloT" | transgene=="7-PoloO")))
```

```
##              numDF denDF   F-value p-value
## (Intercept)     1    95 16.917214  0.0001
## transgene       1    11  0.278728  0.6080
```

#### Inter-species comparisons—test effect on the total number of progeny

##### Do Dpse-polo and poloO differentially affect the total number of progeny?

```
nlme_count_strain(
  subset(bamCounts, sex==" total"
        & (transgene=="4-GA11545" | transgene=="7-PoloO")))
```

```
##              numDF denDF  F-value p-value
## (Intercept)     1   107 6.510926  0.0121
## transgene       1    12 1.954019  0.1875
```

##### Do Dpse-polo and poloT differentially affect the total number of progeny?

```
nlme_count_strain(
  subset(bamCounts, sex==" total"
        & (transgene=="4-GA11545" | transgene=="6-PoloT")))
```

```
##              numDF denDF   F-value p-value
## (Intercept)     1   110 24.471463  <.0001
## transgene       1    12  0.030996  0.8632
```

##### Do Dpse-polo-dup1 and poloO differentially affect the total number of progeny?

```
nlme_count_strain(
  subset(bamCounts, sex==" total"
        & (transgene=="5-GA25172" | transgene=="7-PoloO")))
```

```
##              numDF denDF  F-value p-value
## (Intercept)     1    66 6.427740  0.0136
## transgene       1     6 3.640782  0.1050
```

**Do Dpse-polo-dup1 and poloT differentially affect the total number of progeny?**

```
nlme_count_strain(
  subset(bamCounts, sex==" total"
        & (transgene=="5-GA25172" | transgene=="6-PoloT")))
```

```
##             numDF denDF   F-value p-value
## (Intercept)     1    69 22.444325  <.0001
## transgene       1     6  0.499317  0.5063
```

#### Inter-species comparisons—test effect on the number of female progeny

**Do Dpse-polo and poloO differentially affect the number of female progeny?**

```
nlme_count_strain(
  subset(bamCounts, sex=="female"
        & (transgene=="4-GA11545" | transgene=="7-PoloO")))
```

```
##             numDF denDF  F-value p-value
## (Intercept)     1   107 6.470541  0.0124
## transgene       1    12 0.736488  0.4076
```

**Do Dpse-polo and poloT differentially affect the number of female progeny?**

```
nlme_count_strain(
  subset(bamCounts, sex=="female"
        & (transgene=="4-GA11545" | transgene=="6-PoloT")))
```

```
##             numDF denDF   F-value p-value
## (Intercept)     1   110 27.190514  <.0001
## transgene       1    12  0.075539  0.7881
```

**Do Dpse-polo-dup1 and poloO differentially affect the number of female progeny?**

```
nlme_count_strain(
  subset(bamCounts, sex=="female"
        & (transgene=="5-GA25172" | transgene=="7-PoloO")))
```

```
##             numDF denDF  F-value p-value
## (Intercept)     1    66 6.368644  0.0140
## transgene       1     6 1.472069  0.2706
```

**Do Dpse-polo-dup1 and poloT differentially affect the number of female progeny?**

```
nlme_count_strain(
  subset(bamCounts, sex=="female"
        & (transgene=="5-GA25172" | transgene=="6-PoloT")))
```

```
##             numDF denDF   F-value p-value
## (Intercept)     1    69 27.671325  <.0001
## transgene       1     6  0.763453  0.4158
```

#### Inter-species comparisons—test effect on the number of male progeny

##### Do Dpse-polo and poloO differentially affect the number of male progeny?

```
nlme_count_strain(
  subset(bamCounts, sex=="male"
        & (transgene=="4-GA11545" | transgene=="7-PoloO")))
```

```
##             numDF denDF  F-value p-value
## (Intercept)     1   107 6.573624  0.0117
## transgene       1    12 3.768654  0.0761
```

##### Do Dpse-polo and poloT differentially affect the number of male progeny?

```
nlme_count_strain(
  subset(bamCounts, sex=="male"
        & (transgene=="4-GA11545" | transgene=="6-PoloT")))
```

```
##             numDF denDF   F-value p-value
## (Intercept)     1   110 20.063683  <.0001
## transgene       1    12  0.415797  0.5312
```

##### Do Dpse-polo-dup1 and poloO differentially affect the number of male progeny?

```
nlme_count_strain(
  subset(bamCounts, sex=="male"
        & (transgene=="5-GA25172" | transgene=="7-PoloO")))
```

```
##             numDF denDF  F-value p-value
## (Intercept)     1    66 6.504461  0.0131
## transgene       1     6 6.404146  0.0446
```

##### Do Dpse-polo-dup1 and poloT differentially affect the number of male progeny?

```
nlme_count_strain(
  subset(bamCounts, sex=="male"
        & (transgene=="5-GA25172" | transgene=="6-PoloT")))
```

```
##             numDF denDF   F-value p-value
## (Intercept)     1    69 17.458619  0.0001
## transgene       1     6  0.241523  0.6406
```

#### Logistic regression to test for progeny vs 0 progeny

##### Create column "progeny'' to record 0 progeny vs >0 progeny

```
bamCounts$progeny <- ifelse(bamCounts$counts==0, 0, 1)
```

##### Function to perform logistic regression

```
glmer_progeny_strain <- function(crossData){
    summary(
        glmer(progeny ~ transgene + (1|date) + (1+strainID),
            data = crossData, family = binomial
        )
    )
}
```

#### Test if effect on any progeny produced

##### Do Dpse-polo and Dpse-polo-dup1 differentially affect if there are progeny?

```
glmer_progeny_strain(
  subset(bamCounts, sex==" total"
         & (transgene=="4-GA11545" | transgene=="5-GA25172")))
```

```
## boundary (singular) fit: see help('isSingular')
```

```
## Generalized linear mixed model fit by maximum likelihood (Laplace
##   Approximation) [glmerMod]
##  Family: binomial  ( logit )
## Formula: progeny ~ transgene + (1 | date) + (1 + strainID)
##    Data: crossData
##
##      AIC      BIC   logLik deviance df.resid
##     97.5    110.1    -43.7     87.5       88
##
## Scaled residuals:
##     Min      1Q  Median      3Q     Max
## -4.6904  0.2132  0.5130  0.5941  0.6614
##
## Random effects:
##  Groups Name        Variance Std.Dev.
##  date   (Intercept) 0        0
## Number of obs: 93, groups:  date, 3
##
## Fixed effects:
##                     Estimate Std. Error z value Pr(>|z|)
## (Intercept)           1.0415     0.4749   2.193   0.0283 *
## transgene5-GA25172   2.0496     1.1274   1.818   0.0691 .
## strainIDm14         -0.2148     0.6564  -0.327   0.7435
## strainIDm7           0.2935     0.6915   0.425   0.6712
## ---
## Signif. codes:  0 '***' 0.001 '**' 0.01 '*' 0.05 '.' 0.1 ' ' 1
##
## Correlation of Fixed Effects:
##             (Intr) t5-GA2 stID14
```

```
## tr5-GA25172 -0.421
## strainIDm14 -0.723  0.305
## strainIDm7  -0.687  0.289  0.497
## optimizer (Nelder_Mead) convergence code: 0 (OK)
## boundary (singular) fit: see help('isSingular')
```

**Do Dmel PoloO and PoloT differentially affect if there are progeny?**

```
glmer_progeny_strain(
  subset(bamCounts, sex==" total"
       & (transgene=="6-PoloT" | transgene=="7-PoloO")))
```

```
## fixed-effect model matrix is rank deficient so dropping 1 column / coefficient
```

```
## boundary (singular) fit: see help('isSingular')
```

```
## Generalized linear mixed model fit by maximum likelihood (Laplace
##   Approximation) [glmerMod]
##  Family: binomial  ( logit )
## Formula: progeny ~ transgene + (1 | date) + (1 + strainID)
##    Data: crossData
##
##      AIC      BIC   logLik deviance df.resid
##    126.6    140.1    -58.3    116.6      106
##
## Scaled residuals:
##     Min      1Q  Median      3Q     Max
## -2.1909  0.4564  0.4879  0.5401  0.6860
##
## Random effects:
##  Groups Name        Variance Std.Dev.
##  date   (Intercept) 0        0
## Number of obs: 111, groups:  date, 4
##
## Fixed effects:
##                  Estimate Std. Error z value Pr(>|z|)
## (Intercept)        1.4351     0.4976   2.884  0.00393 **
## transgene7-PoloO  -0.6813     0.6568  -1.037  0.29962
## strainIDm12        0.8148     0.6523   1.249  0.21159
## strainIDm9        -0.2029     0.6574  -0.309  0.75754
## ---
## Signif. codes:  0 '***' 0.001 '**' 0.01 '*' 0.05 '.' 0.1 ' ' 1
##
## Correlation of Fixed Effects:
##             (Intr) tr7-PO stID12
## trnsgn7-PlO -0.758
## strainIDm12  0.000 -0.429
## strainIDm9  -0.757  0.573  0.000
## fit warnings:
## fixed-effect model matrix is rank deficient so dropping 1 column / coefficient
## optimizer (Nelder_Mead) convergence code: 0 (OK)
## boundary (singular) fit: see help('isSingular')
```

#### Test if effect on if female progeny are produced?

##### Do Dpse-polo and Dpse-polo-dup1 differentially affect if there are female progeny?

```
glmer_progeny_strain(
  subset(bamCounts, sex=="female"
       & (transgene=="4-GA11545" | transgene=="5-GA25172")))
```

```
## boundary (singular) fit: see help('isSingular')
```

```
## Generalized linear mixed model fit by maximum likelihood (Laplace
##   Approximation) [glmerMod]
##  Family: binomial  ( logit )
## Formula: progeny ~ transgene + (1 | date) + (1 + strainID)
##    Data: crossData
##
##      AIC      BIC   logLik deviance df.resid
##    111.8    124.5    -50.9    101.8       88
##
## Scaled residuals:
##     Min      1Q  Median      3Q     Max
## -2.5820 -1.3693  0.5130  0.7303  0.7303
##
## Random effects:
##  Groups Name        Variance Std.Dev.
##  date   (Intercept) 0        0
## Number of obs: 93, groups:  date, 3
##
## Fixed effects:
##                     Estimate Std. Error z value Pr(>|z|)
## (Intercept)         6.286e-01  4.378e-01   1.436   0.1510
## transgene5-GA25172  1.269e+00  7.583e-01   1.673   0.0944 .
## strainIDm14        -9.740e-16  6.191e-01   0.000   1.0000
## strainIDm7          7.064e-01  6.666e-01   1.060   0.2893
## ---
## Signif. codes:  0 '***' 0.001 '**' 0.01 '*' 0.05 '.' 0.1 ' ' 1
##
## Correlation of Fixed Effects:
##             (Intr) t5-GA2 stID14
## tr5-GA25172 -0.577
## strainIDm14 -0.707  0.408
## strainIDm7  -0.657  0.379  0.464
## optimizer (Nelder_Mead) convergence code: 0 (OK)
## boundary (singular) fit: see help('isSingular')
```

##### Do Dmel PoloO and PoloT differentially affect if there are female progeny?

```
glmer_progeny_strain(
  subset(bamCounts, sex=="female"
       & (transgene=="6-PoloT" | transgene=="7-PoloO")))
```

```
## fixed-effect model matrix is rank deficient so dropping 1 column / coefficient
```

```
## boundary (singular) fit: see help('isSingular')
```

```
## Generalized linear mixed model fit by maximum likelihood (Laplace
```

```
##   Approximation) [glmerMod]
##  Family: binomial  ( logit )
## Formula: progeny ~ transgene + (1 | date) + (1 + strainID)
##    Data: crossData
##
##      AIC      BIC   logLik deviance df.resid
##    145.6    159.2    -67.8    135.6      106
##
## Scaled residuals:
##     Min      1Q  Median      3Q     Max
## -1.7728 -1.3333  0.5641  0.7276  0.7500
##
## Random effects:
##  Groups Name        Variance Std.Dev.
##  date   (Intercept) 0        0
## Number of obs: 111, groups:  date, 4
##
## Fixed effects:
##                   Estimate Std. Error z value Pr(>|z|)
## (Intercept)        0.63599    0.41223   1.543    0.123
## transgene7-PoloO -0.06062    0.58613  -0.103    0.918
## strainIDm12        0.56977    0.60160   0.947    0.344
## strainIDm9         0.25783    0.57140   0.451    0.652
##
## Correlation of Fixed Effects:
##             (Intr) tr7-PO stID12
## trnsgn7-PlO -0.703
## strainIDm12  0.000 -0.492
## strainIDm9  -0.721  0.507  0.000
## fit warnings:
## fixed-effect model matrix is rank deficient so dropping 1 column / coefficient
## optimizer (Nelder_Mead) convergence code: 0 (OK)
## boundary (singular) fit: see help('isSingular')
```

#### Test if effect on if male progeny are produced?

##### Do Dpse-polo and Dpse-polo-dup1 differentially affect if there are male progeny?

```r
glmer_progeny_strain(
  subset(bamCounts, sex=="male"
        & (transgene=="4-GA11545" | transgene=="5-GA25172")))
```

```
## boundary (singular) fit: see help('isSingular')
```

```
## Generalized linear mixed model fit by maximum likelihood (Laplace
##   Approximation) [glmerMod]
##  Family: binomial  ( logit )
## Formula: progeny ~ transgene + (1 | date) + (1 + strainID)
##    Data: crossData
##
##      AIC      BIC   logLik deviance df.resid
##     99.9    112.6    -44.9     89.9       88
##
## Scaled residuals:
##     Min      1Q  Median      3Q     Max
```

```
## -4.6904  0.2132  0.5774  0.5941  0.6614
##
## Random effects:
##  Groups Name        Variance  Std.Dev.
##  date   (Intercept) 1.631e-16 1.277e-08
## Number of obs: 93, groups:  date, 3
##
## Fixed effects:
##                     Estimate Std. Error z value Pr(>|z|)
## (Intercept)          1.04145    0.47486   2.193   0.0283 *
## transgene5-GA25172  2.04959    1.12736   1.818   0.0691 .
## strainIDm14        -0.21478    0.65639  -0.327   0.7435
## strainIDm7          0.05716    0.66911   0.085   0.9319
## ---
## Signif. codes:  0 '***' 0.001 '**' 0.01 '*' 0.05 '.' 0.1 ' ' 1
##
## Correlation of Fixed Effects:
##             (Intr) t5-GA2 stID14
## tr5-GA25172 -0.421
## strainIDm14 -0.723  0.305
## strainIDm7  -0.710  0.299  0.513
## optimizer (Nelder_Mead) convergence code: 0 (OK)
## boundary (singular) fit: see help('isSingular')
```

##### Do Dmel PoloO and PoloT differentially affect if there are male progeny?

```
glmer_progeny_strain(
  subset(bamCounts, sex=="male"
       & (transgene=="6-PoloT" | transgene=="7-PoloO")))
```

```
## fixed-effect model matrix is rank deficient so dropping 1 column / coefficient
```

```
## Generalized linear mixed model fit by maximum likelihood (Laplace
##   Approximation) [glmerMod]
##  Family: binomial  ( logit )
## Formula: progeny ~ transgene + (1 | date) + (1 + strainID)
##    Data: crossData
##
##      AIC      BIC   logLik deviance df.resid
##    142.3    155.8    -66.1    132.3      106
##
## Scaled residuals:
##     Min      1Q  Median      3Q     Max
## -2.2837 -1.1639  0.5080  0.6775  0.8614
##
## Random effects:
##  Groups Name        Variance Std.Dev.
##  date   (Intercept) 0.1099   0.3315
## Number of obs: 111, groups:  date, 4
##
## Fixed effects:
##                   Estimate Std. Error z value Pr(>|z|)
## (Intercept)         1.5589     0.5753   2.710  0.00673 **
## transgene7-PoloO  -1.0513     0.6496  -1.618  0.10559
```

```
## strainIDm12        0.7158     0.6031   1.187  0.23527  
## strainIDm9        -1.0564     0.6419  -1.646  0.09982 .
## ---
## Signif. codes:  0 '***' 0.001 '**' 0.01 '*' 0.05 '.' 0.1 ' ' 1
##
## Correlation of Fixed Effects:
##             (Intr) tr7-PO stID12
## trnsgn7-PlO -0.694              
## strainIDm12 -0.025 -0.431       
## strainIDm9  -0.769  0.614  0.017
## fit warnings:
## fixed-effect model matrix is rank deficient so dropping 1 column / coefficient
```

#### Inter-species comparison of progeny vs no progeny

##### Do Dpse-polo and poloO differentially affect if progeny produced?

```
glmer_progeny_strain(
  subset(bamCounts, sex==" total"
       & (transgene=="4-GA11545" | transgene=="7-PoloO")))
```

```
## fixed-effect model matrix is rank deficient so dropping 1 column / coefficient
```

```
## boundary (singular) fit: see help('isSingular')
```

```
## Generalized linear mixed model fit by maximum likelihood (Laplace
##   Approximation) [glmerMod]
##  Family: binomial  ( logit )
## Formula: progeny ~ transgene + (1 | date) + (1 + strainID)
##    Data: crossData
##
##      AIC      BIC   logLik deviance df.resid 
##    149.2    166.2    -68.6    137.2      118 
##
## Scaled residuals: 
##      Min       1Q   Median       3Q      Max 
## -2.19089 -0.02211  0.51299  0.66144  0.68599 
##
## Random effects:
##  Groups Name        Variance Std.Dev.
##  date   (Intercept) 0        0       
## Number of obs: 124, groups:  date, 4
##
## Fixed effects:
##                   Estimate Std. Error z value Pr(>|z|)  
## (Intercept)         1.0415     0.4749   2.193   0.0283 *
## transgene7-PoloO  -0.2877     0.6398  -0.450   0.6530  
## strainIDm12         0.8148     0.6523   1.249   0.2116  
## strainIDm14       -0.2148     0.6564  -0.327   0.7435  
## strainIDm7          0.2935     0.6915   0.425   0.6712  
## ---
## Signif. codes:  0 '***' 0.001 '**' 0.01 '*' 0.05 '.' 0.1 ' ' 1
##
## Correlation of Fixed Effects:
##             (Intr) tr7-PO stID12 stID14
```

```
## trnsgn7-PlO -0.742
## strainIDm12  0.000 -0.440
## strainIDm14 -0.723  0.537  0.000
## strainIDm7  -0.687  0.510  0.000  0.497
## fit warnings:
## fixed-effect model matrix is rank deficient so dropping 1 column / coefficient
## optimizer (Nelder_Mead) convergence code: 0 (OK)
## boundary (singular) fit: see help('isSingular')
```

##### Do Dpse-polo and poloT differentially affect if progeny produced?

```
glmer_progeny_strain(
  subset(bamCounts, sex==" total"
        & (transgene=="4-GA11545" | transgene=="6-PoloT")))
```

```
## fixed-effect model matrix is rank deficient so dropping 1 column / coefficient
```

```
## boundary (singular) fit: see help('isSingular')
```

```
## Generalized linear mixed model fit by maximum likelihood (Laplace
##   Approximation) [glmerMod]
##  Family: binomial  ( logit )
## Formula: progeny ~ transgene + (1 | date) + (1 + strainID)
##    Data: crossData
##
##      AIC      BIC   logLik deviance df.resid
##    149.8    166.9    -68.9    137.8      121
##
## Scaled residuals:
##     Min      1Q  Median      3Q     Max
## -2.0494  0.4879  0.5130  0.5941  0.6614
##
## Random effects:
##  Groups Name        Variance Std.Dev.
##  date   (Intercept) 0        0
## Number of obs: 127, groups:  date, 4
##
## Fixed effects:
##                   Estimate Std. Error z value Pr(>|z|)
## (Intercept)         1.0415     0.4749   2.193   0.0283 *
## transgene6-PoloT    0.1907     0.6403   0.298   0.7659
## strainIDf15         0.2029     0.6574   0.309   0.7575
## strainIDm14        -0.2148     0.6564  -0.327   0.7435
## strainIDm7          0.2935     0.6915   0.425   0.6712
## ---
## Signif. codes:  0 '***' 0.001 '**' 0.01 '*' 0.05 '.' 0.1 ' ' 1
##
## Correlation of Fixed Effects:
##             (Intr) tr6-PT stID15 stID14
## trnsgn6-PlT -0.742
## strainIDf15  0.000 -0.438
## strainIDm14 -0.723  0.536  0.000
## strainIDm7  -0.687  0.509  0.000  0.497
## fit warnings:
## fixed-effect model matrix is rank deficient so dropping 1 column / coefficient
```

```
## optimizer (Nelder_Mead) convergence code: 0 (OK)
## boundary (singular) fit: see help('isSingular')
```

**Do Dpse-polo-dup1 and poloO differentially affect if progeny produced?**

```r
glmer_progeny_strain(
  subset(bamCounts, sex==" total"
         & (transgene=="5-GA25172" | transgene=="7-PoloO")))
```

```
## fixed-effect model matrix is rank deficient so dropping 1 column / coefficient
```

```
## boundary (singular) fit: see help('isSingular')
```

```
## Generalized linear mixed model fit by maximum likelihood (Laplace
##   Approximation) [glmerMod]
##  Family: binomial  ( logit )
## Formula: progeny ~ transgene + (1 | date) + (1 + strainID)
##    Data: crossData
##
##      AIC      BIC   logLik deviance df.resid
##     74.2     83.6    -33.1     66.2       73
##
## Scaled residuals:
##     Min      1Q  Median      3Q     Max
## -4.6904  0.2132  0.4564  0.4564  0.6860
##
## Random effects:
##  Groups Name        Variance Std.Dev.
##  date   (Intercept) 0        0
## Number of obs: 77, groups:  date, 4
##
## Fixed effects:
##                  Estimate Std. Error z value Pr(>|z|)
## (Intercept)        3.0910     1.0225   3.023   0.0025 **
## transgene7-PoloO  -2.3373     1.1087  -2.108   0.0350 *
## strainIDm12        0.8148     0.6523   1.249   0.2116
## ---
## Signif. codes:  0 '***' 0.001 '**' 0.01 '*' 0.05 '.' 0.1 ' ' 1
##
## Correlation of Fixed Effects:
##             (Intr) tr7-PO
## trnsgn7-PlO -0.922
## strainIDm12  0.000 -0.254
## fit warnings:
## fixed-effect model matrix is rank deficient so dropping 1 column / coefficient
## optimizer (Nelder_Mead) convergence code: 0 (OK)
## boundary (singular) fit: see help('isSingular')
```

**Do Dpse-polo-dup1 and poloT differentially affect if progeny produced?**

```r
glmer_progeny_strain(
  subset(bamCounts, sex==" total"
         & (transgene=="5-GA25172" | transgene=="6-PoloT")))
```

```
## fixed-effect model matrix is rank deficient so dropping 1 column / coefficient
```

```
## Generalized linear mixed model fit by maximum likelihood (Laplace
##   Approximation) [glmerMod]
##  Family: binomial  ( logit )
## Formula: progeny ~ transgene + (1 | date) + (1 + strainID)
##    Data: crossData
##
##      AIC      BIC   logLik deviance df.resid
##     74.8     84.3    -33.4     66.8       76
##
## Scaled residuals:
##     Min      1Q  Median      3Q     Max
## -4.5855  0.2064  0.4722  0.5242  0.5590
##
## Random effects:
##  Groups Name        Variance Std.Dev.
##  date   (Intercept) 0.03423  0.185
## Number of obs: 80, groups:  date, 4
##
## Fixed effects:
##                  Estimate Std. Error z value Pr(>|z|)
## (Intercept)        3.1371     1.0873   2.885  0.00391 **
## transgene6-PoloT  -1.8825     1.1253  -1.673  0.09434 .
## strainIDf15        0.2140     0.6653   0.322  0.74773
## ---
## Signif. codes:  0 '***' 0.001 '**' 0.01 '*' 0.05 '.' 0.1 ' ' 1
##
## Correlation of Fixed Effects:
##             (Intr) tr6-PT
## trnsgn6-PlT -0.909
## strainIDf15  0.043 -0.270
## fit warnings:
## fixed-effect model matrix is rank deficient so dropping 1 column / coefficient
```

#### Inter-species comparison of female progeny vs no female progeny

##### Do Dpse-polo and poloO differentially affect if female progeny produced?

```
glmer_progeny_strain(
  subset(bamCounts, sex=="female"
        & (transgene=="4-GA11545" | transgene=="7-PoloO")))
```

```
## fixed-effect model matrix is rank deficient so dropping 1 column / coefficient
```

```
## boundary (singular) fit: see help('isSingular')
```

```
## Generalized linear mixed model fit by maximum likelihood (Laplace
##   Approximation) [glmerMod]
##  Family: binomial  ( logit )
## Formula: progeny ~ transgene + (1 | date) + (1 + strainID)
##    Data: crossData
##
##      AIC      BIC   logLik deviance df.resid
##    160.7    177.7    -74.4    148.7      118
##
## Scaled residuals:
```

```
##     Min      1Q  Median      3Q     Max 
## -1.9494 -1.3333  0.5641  0.7303  0.7500 
## 
## Random effects:
##  Groups Name        Variance Std.Dev.
##  date   (Intercept) 0        0       
## Number of obs: 124, groups:  date, 4
## 
## Fixed effects:
##                     Estimate Std. Error z value Pr(>|z|)
## (Intercept)        6.286e-01  4.378e-01   1.436    0.151
## transgene7-PoloO -5.324e-02  6.044e-01  -0.088    0.930
## strainIDm12        5.698e-01  6.016e-01   0.947    0.344
## strainIDm14        3.044e-16  6.191e-01   0.000    1.000
## strainIDm7         7.064e-01  6.666e-01   1.060    0.289
## 
## Correlation of Fixed Effects:
##             (Intr) tr7-PO stID12 stID14
## trnsgn7-PlO -0.724                     
## strainIDm12  0.000 -0.477              
## strainIDm14 -0.707  0.512  0.000       
## strainIDm7  -0.657  0.476  0.000  0.464
## fit warnings:
## fixed-effect model matrix is rank deficient so dropping 1 column / coefficient
## optimizer (Nelder_Mead) convergence code: 0 (OK)
## boundary (singular) fit: see help('isSingular')
```

##### Do Dpse-polo and poloT differentially affect if female progeny produced?

```r
glmer_progeny_strain(
  subset(bamCounts, sex=="female"
       & (transgene=="4-GA11545" | transgene=="6-PoloT")))
```

```
## fixed-effect model matrix is rank deficient so dropping 1 column / coefficient

## boundary (singular) fit: see help('isSingular')

## Generalized linear mixed model fit by maximum likelihood (Laplace
##   Approximation) [glmerMod]
##  Family: binomial  ( logit )
## Formula: progeny ~ transgene + (1 | date) + (1 + strainID)
##    Data: crossData
## 
##      AIC      BIC   logLik deviance df.resid 
##    166.9    184.0    -77.4    154.9      121 
## 
## Scaled residuals: 
##     Min      1Q  Median      3Q     Max 
## -1.9494 -1.3693  0.6396  0.7276  0.7303 
## 
## Random effects:
##  Groups Name        Variance Std.Dev.
##  date   (Intercept) 0        0       
## Number of obs: 127, groups:  date, 4
## 
```

```
## Fixed effects:
##                    Estimate Std. Error z value Pr(>|z|)
## (Intercept)       6.286e-01  4.378e-01   1.436    0.151
## transgene6-PoloT  2.652e-01  5.901e-01   0.449    0.653
## strainIDf15      -2.578e-01  5.714e-01  -0.451    0.652
## strainIDm14       3.957e-15  6.191e-01   0.000    1.000
## strainIDm7        7.064e-01  6.666e-01   1.060    0.289
##
## Correlation of Fixed Effects:
##             (Intr) tr6-PT stID15 stID14
## trnsgn6-PlT -0.742
## strainIDf15  0.000 -0.464
## strainIDm14 -0.707  0.525  0.000
## strainIDm7  -0.657  0.487  0.000  0.464
## fit warnings:
## fixed-effect model matrix is rank deficient so dropping 1 column / coefficient
## optimizer (Nelder_Mead) convergence code: 0 (OK)
## boundary (singular) fit: see help('isSingular')
```

##### Do Dpse-polo-dup1 and poloO differentially affect if female progeny produced?

```r
glmer_progeny_strain(
  subset(bamCounts, sex=="female"
        & (transgene=="5-GA25172" | transgene=="7-PoloO")))
```

```
## fixed-effect model matrix is rank deficient so dropping 1 column / coefficient
```

```
## Generalized linear mixed model fit by maximum likelihood (Laplace
##   Approximation) [glmerMod]
##  Family: binomial  ( logit )
## Formula: progeny ~ transgene + (1 | date) + (1 + strainID)
##    Data: crossData
##
##      AIC      BIC   logLik deviance df.resid
##     90.5     99.8    -41.2     82.5       73
##
## Scaled residuals:
##     Min      1Q  Median      3Q     Max
## -2.7083  0.3642  0.4139  0.6111  0.8148
##
## Random effects:
##  Groups Name        Variance Std.Dev.
##  date   (Intercept) 0.06526  0.2555
## Number of obs: 77, groups:  date, 4
##
## Fixed effects:
##                  Estimate Std. Error z value Pr(>|z|)
## (Intercept)        1.9507     0.6728   2.899  0.00374 **
## transgene7-PoloO  -1.3544     0.7597  -1.783  0.07463 .
## strainIDm12        0.5753     0.6067   0.948  0.34301
## ---
## Signif. codes:  0 '***' 0.001 '**' 0.01 '*' 0.05 '.' 0.1 ' ' 1
##
## Correlation of Fixed Effects:
```

```
##              (Intr) tr7-PO
## trnsgn7-PlO -0.808
## strainIDm12  0.005 -0.386
## fit warnings:
## fixed-effect model matrix is rank deficient so dropping 1 column / coefficient
```

**Do Dpse-polo-dup1 and poloT differentially affect if female progeny produced?**

```
glmer_progeny_strain(
  subset(bamCounts, sex=="female"
        & (transgene=="5-GA25172" | transgene=="6-PoloT")))
```

```
## fixed-effect model matrix is rank deficient so dropping 1 column / coefficient
```

```
## boundary (singular) fit: see help('isSingular')
```

```
## Generalized linear mixed model fit by maximum likelihood (Laplace
##   Approximation) [glmerMod]
##  Family: binomial  ( logit )
## Formula: progeny ~ transgene + (1 | date) + (1 + strainID)
##    Data: crossData
##
##      AIC      BIC   logLik deviance df.resid
##     96.7    106.2    -44.4     88.7       76
##
## Scaled residuals:
##     Min      1Q  Median      3Q     Max
## -2.5820 -1.3744  0.3873  0.6396  0.7276
##
## Random effects:
##  Groups Name        Variance Std.Dev.
##  date   (Intercept) 0        0
## Number of obs: 80, groups:  date, 4
##
## Fixed effects:
##                   Estimate Std. Error z value Pr(>|z|)
## (Intercept)         1.8971     0.6191   3.064  0.00218 **
## transgene6-PoloT  -1.0033     0.7348  -1.365  0.17211
## strainIDf15       -0.2578     0.5714  -0.451  0.65183
## ---
## Signif. codes:  0 '***' 0.001 '**' 0.01 '*' 0.05 '.' 0.1 ' ' 1
##
## Correlation of Fixed Effects:
##             (Intr) tr6-PT
## trnsgn6-PlT -0.843
## strainIDf15  0.000 -0.373
## fit warnings:
## fixed-effect model matrix is rank deficient so dropping 1 column / coefficient
## optimizer (Nelder_Mead) convergence code: 0 (OK)
## boundary (singular) fit: see help('isSingular')
```

#### Inter-species comparison of male progeny vs no male progeny

##### Do Dpse-polo and poloO differentially affect if male progeny produced?

```
glmer_progeny_strain(
  subset(bamCounts, sex=="male"
       & (transgene=="4-GA11545" | transgene=="7-PoloO")))
```

```
## fixed-effect model matrix is rank deficient so dropping 1 column / coefficient

## boundary (singular) fit: see help('isSingular')

## Generalized linear mixed model fit by maximum likelihood (Laplace
##   Approximation) [glmerMod]
##  Family: binomial  ( logit )
## Formula: progeny ~ transgene + (1 | date) + (1 + strainID)
##    Data: crossData
##
##      AIC      BIC   logLik deviance df.resid
##    159.4    176.3    -73.7    147.4      118
##
## Scaled residuals:
##     Min      1Q  Median      3Q     Max
## -1.7728 -1.2247  0.5774  0.6109  0.8165
##
## Random effects:
##  Groups Name        Variance Std.Dev.
##  date   (Intercept) 0        0
## Number of obs: 124, groups:  date, 4
##
## Fixed effects:
##                  Estimate Std. Error z value Pr(>|z|)
## (Intercept)       1.04145    0.47486   2.193   0.0283 *
## transgene7-PoloO -0.63599    0.62622  -1.016   0.3098
## strainIDm12       0.73967    0.59580   1.241   0.2144
## strainIDm14      -0.21478    0.65639  -0.327   0.7435
## strainIDm7        0.05716    0.66911   0.085   0.9319
## ---
## Signif. codes:  0 '***' 0.001 '**' 0.01 '*' 0.05 '.' 0.1 ' ' 1
##
## Correlation of Fixed Effects:
##             (Intr) tr7-PO stID12 stID14
## trnsgn7-PlO -0.758
## strainIDm12  0.000 -0.447
## strainIDm14 -0.723  0.549  0.000
## strainIDm7  -0.710  0.538  0.000  0.513
## fit warnings:
## fixed-effect model matrix is rank deficient so dropping 1 column / coefficient
## optimizer (Nelder_Mead) convergence code: 0 (OK)
## boundary (singular) fit: see help('isSingular')
```

**Do Dpse-polo and poloT differentially affect if male progeny produced?**

```r
glmer_progeny_strain(
  subset(bamCounts, sex=="male"
       & (transgene=="4-GA11545" | transgene=="6-PoloT")))
```

```
## fixed-effect model matrix is rank deficient so dropping 1 column / coefficient

## boundary (singular) fit: see help('isSingular')

## Generalized linear mixed model fit by maximum likelihood (Laplace
##   Approximation) [glmerMod]
##  Family: binomial  ( logit )
## Formula: progeny ~ transgene + (1 | date) + (1 + strainID)
##    Data: crossData
##
##      AIC      BIC   logLik deviance df.resid
##    160.5    177.6    -74.2    148.5      121
##
## Scaled residuals:
##     Min      1Q  Median      3Q     Max
## -2.0494 -1.2583  0.5774  0.6614  0.7947
##
## Random effects:
##  Groups Name        Variance Std.Dev.
##  date   (Intercept) 0        0
## Number of obs: 127, groups:  date, 4
##
## Fixed effects:
##                   Estimate Std. Error z value Pr(>|z|)
## (Intercept)        1.04145    0.47486   2.193   0.0283 *
## transgene6-PoloT -0.58192    0.60121  -0.968   0.3331
## strainIDf15        0.97555    0.61934   1.575   0.1152
## strainIDm14       -0.21478    0.65639  -0.327   0.7435
## strainIDm7         0.05716    0.66911   0.085   0.9319
## ---
## Signif. codes:  0 '***' 0.001 '**' 0.01 '*' 0.05 '.' 0.1 ' ' 1
##
## Correlation of Fixed Effects:
##             (Intr) tr6-PT stID15 stID14
## trnsgn6-PlT -0.790
## strainIDf15  0.000 -0.365
## strainIDm14 -0.723  0.571  0.000
## strainIDm7  -0.710  0.561  0.000  0.513
## fit warnings:
## fixed-effect model matrix is rank deficient so dropping 1 column / coefficient
## optimizer (Nelder_Mead) convergence code: 0 (OK)
## boundary (singular) fit: see help('isSingular')
```

##### Do Dpse-polo-dup1 and poloO differentially affect if male progeny produced?

```r
glmer_progeny_strain(
  subset(bamCounts, sex=="male"
       & (transgene=="5-GA25172" | transgene=="7-PoloO")))
```

```
## fixed-effect model matrix is rank deficient so dropping 1 column / coefficient
```

```
## boundary (singular) fit: see help('isSingular')
```

```
## Generalized linear mixed model fit by maximum likelihood (Laplace
##   Approximation) [glmerMod]
##  Family: binomial  ( logit )
## Formula: progeny ~ transgene + (1 | date) + (1 + strainID)
##    Data: crossData
##
##      AIC      BIC   logLik deviance df.resid
##     81.9     91.3    -37.0     73.9       73
##
## Scaled residuals:
##     Min      1Q  Median      3Q     Max
## -4.6904  0.2132  0.2132  0.5641  0.8165
##
## Random effects:
##  Groups Name        Variance Std.Dev.
##  date   (Intercept) 0        0
## Number of obs: 77, groups:  date, 4
##
## Fixed effects:
##                  Estimate Std. Error z value Pr(>|z|)
## (Intercept)        3.0910     1.0225   3.023   0.0025 **
## transgene7-PoloO  -2.6856     1.1010  -2.439   0.0147 *
## strainIDm12        0.7397     0.5958   1.241   0.2144
## ---
## Signif. codes:  0 '***' 0.001 '**' 0.01 '*' 0.05 '.' 0.1 ' ' 1
##
## Correlation of Fixed Effects:
##             (Intr) tr7-PO
## trnsgn7-PlO -0.929
## strainIDm12  0.000 -0.254
## fit warnings:
## fixed-effect model matrix is rank deficient so dropping 1 column / coefficient
## optimizer (Nelder_Mead) convergence code: 0 (OK)
## boundary (singular) fit: see help('isSingular')
```

##### Do Dpse-polo-dup1 and poloT differentially affect if male progeny produced?

```r
glmer_progeny_strain(
  subset(bamCounts, sex=="male"
       & (transgene=="5-GA25172" | transgene=="6-PoloT")))
```

```
## fixed-effect model matrix is rank deficient so dropping 1 column / coefficient
```

```
## Generalized linear mixed model fit by maximum likelihood (Laplace
##   Approximation) [glmerMod]
##  Family: binomial  ( logit )
```

```
## Formula: progeny ~ transgene + (1 | date) + (1 + strainID)
##    Data: crossData
##
##      AIC      BIC   logLik deviance df.resid
##     83.0     92.5    -37.5     75.0       76
##
## Scaled residuals:
##     Min      1Q  Median      3Q     Max
## -4.5082  0.1928  0.3347  0.5148  0.8534
##
## Random effects:
##  Groups Name        Variance Std.Dev.
##  date   (Intercept) 0.07105  0.2666
## Number of obs: 80, groups:  date, 4
##
## Fixed effects:
##                  Estimate Std. Error z value Pr(>|z|)
## (Intercept)        3.1926     1.0980   2.908  0.00364 **
## transgene6-PoloT  -2.6947     1.1141  -2.419  0.01557 *
## strainIDf15        1.0109     0.6383   1.584  0.11326
## ---
## Signif. codes:  0 '***' 0.001 '**' 0.01 '*' 0.05 '.' 0.1 ' ' 1
##
## Correlation of Fixed Effects:
##             (Intr) tr6-PT
## trnsgn6-PlT -0.928
## strainIDf15  0.073 -0.241
## fit warnings:
## fixed-effect model matrix is rank deficient so dropping 1 column / coefficient
```

#### Plot the number of male, female, and total progeny

##### Prepare the data

```
bamCounts$Xaxis <- NA
bamCounts$Xaxis[bamCounts$transgene=="4-GA11545"] <- "Dpse-polo"
bamCounts$Xaxis[bamCounts$transgene=="5-GA25172"] <- "Dpse-polo-dup1"
bamCounts$Xaxis[bamCounts$transgene=="6-PoloT"] <- " polo (testis)"
bamCounts$Xaxis[bamCounts$transgene=="7-PoloO"] <- " polo (ovary)"
```

##### Make graph

```
ggplot(bamCounts, aes(y=counts, x=Xaxis)) +
    geom_boxplot(outlier.shape = NA) +
    geom_jitter(size=0.5, width=0.2) +
    facet_wrap(~sex, ncol=1, scales="free_y") +
    scale_y_continuous(name="number of progeny") +
    scale_x_discrete(name="") +
    theme_bw() +
    theme(axis.text.x = element_text(angle = 45, vjust = 1, hjust=1))
```

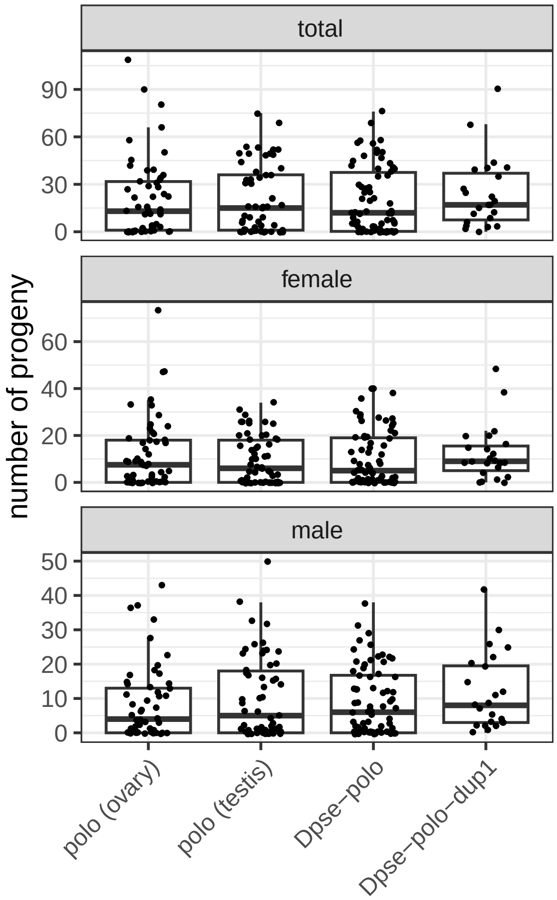


#### What is the effect of each transgene on the relative frequency of males and females?

##### Prepare data for analysis

```
bam.df <- data.frame(
    date = bams$Cross.date,
    vial = bams$vial,
    strain = bams$strain,
    counts = c(bams$males, bams$females),
    sex = c(rep("male",length(bams$males)), rep("female",length(bams$females)))
)
bam.df$vialID <- paste(bam.df$date, bam.df$vial, sep="_")
bam.df <- bam.df %>% separate(
  strain, into=c("transgene", "strainID"), sep=" ", remove=FALSE)
```

##### Create function to perform statistical tests

```
nlme_sex_strain <- function(crossData){
    anova(
        lme(fixed = counts ~ sex + vialID,
            data = crossData,
            random = list(~1|date, ~1|strainID)
        )
```

```
    )
}
```

**Does expression of Dpse-polo in the D. melanogaster male germline affect the relative numbers of male and female progeny sired?**

```
nlme_sex_strain(subset(bam.df, transgene=="4-GA11545")) # Dpse-polo
```

```
##             numDF denDF  F-value p-value
## (Intercept)     1    61 68.90748  <.0001
## sex             1    61  3.50099  0.0661
## vialID         69    61 13.73310  <.0001
```

**Does expression of Dpse-polo-dup1 in the D. melanogaster male germline affect the relative numbers of male and female progeny sired?**

```
nlme_sex_strain(subset(bam.df, transgene=="5-GA25172")) # Dpse-polo-dup1
```

```
##             numDF denDF  F-value p-value
## (Intercept)     1    20 33.52007  <.0001
## sex             1    20  0.06745  0.7977
## vialID         22    20 10.67630  <.0001
```

**Does expression of PoloT in the D. melanogaster male germline affect the relative numbers of male and female progeny sired?**

```
nlme_sex_strain(subset(bam.df, transgene=="6-PoloT"))
```

```
##             numDF denDF  F-value p-value
## (Intercept)     1    49 42.90222  <.0001
## sex             1    49  0.17536  0.6772
## vialID         56    49  9.99859  <.0001
```

**Does expression of PoloO in the D. melanogaster male germline affect the relative numbers of male and female progeny sired?**

```
nlme_sex_strain(subset(bam.df, transgene=="7-PoloO"))
```

```
##             numDF denDF   F-value p-value
## (Intercept)     1    46 30.625273  <.0001
## sex             1    46  9.348539  0.0037
## vialID         53    46  5.669052  <.0001
```

#### Plot proportion of females for each transgene

##### Prepare data for plotting

```
bams <- bams %>% separate(
  strain, into=c("transgene", "strainID"), sep=" ", remove=FALSE)
bams$Xaxis <- NA
bams$Xaxis[bams$transgene=="4-GA11545"] <- "Dpse-polo"
bams$Xaxis[bams$transgene=="5-GA25172"] <- "Dpse-polo-dup1"
bams$Xaxis[bams$transgene=="6-PoloT"] <- " polo (testis)"
```

```
bams$Xaxis[bams$transgene=="7-Polo0"] <- " polo (ovary)"
bams$freqFem <- bams$females/(bams$females + bams$males)
```

##### Create graph

```
ggplot(bams, aes(y=freqFem, x=Xaxis)) +
    geom_boxplot(outlier.shape=NA) +
    geom_jitter(size=0.5, width=0.2) +
    scale_y_continuous(name="frequency females") +
    scale_x_discrete(name="") +
    geom_hline(yintercept=0.5, linetype=2) +
    theme_bw() +
    theme(axis.text.x = element_text(angle = 45, vjust = 1, hjust=1))
```

```
## Warning: Removed 44 rows containing non-finite values (`stat_boxplot()`).
```

```
## Warning: Removed 44 rows containing missing values (`geom_point()`).
```

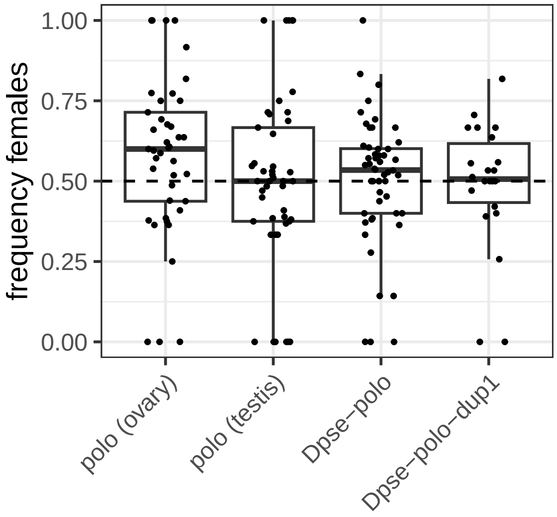
